# Carbon ion irradiation plus CTLA4 blockade elicits therapeutic immune responses in a murine tumor model

**DOI:** 10.1101/2022.07.22.500608

**Authors:** Laura Hartmann, Wolfram Osen, Oliver L. Eichmüller, Theresa Kordaß, Jennifer Furkel, Elke Dickes, Carissa Reid, Jürgen Debus, Stephan Brons, Amir Abdollahi, Mahmoud Moustafa, Stefan Rieken, Stefan B. Eichmüller

**Affiliations:** Research Group GMP & T Cell Therapy, German Cancer Research Center (DKFZ), Heidelberg, Germany; Faculty of Biosciences, Heidelberg University, Heidelberg, Germany; Institute of Molecular Biotechnology (IMBA), Austrian Academy of Sciences, Vienna Biocenter (VBC), Vienna, Austria; National Center for Tumor Diseases (NCT), Heidelberg University Hospital (UKHD) and German Cancer Research Center, Heidelberg, Germany; CCU Translational Radiation Oncology, German Cancer Consortium (DKTK) Core-Center Heidelberg, Heidelberg University Hospital (UKHD) and German Cancer Research Center (DKFZ), Heidelberg, Germany; Heidelberg Institute of Radiation Oncology (HIRO), National Center for Radiation Oncology (NCRO), Heidelberg, Germany; Heidelberg Ion-Beam Therapy Center (HIT), Department of Radiation Oncology, Heidelberg University Hospital (UKHD), Heidelberg, Germany; Division of Biostatistics, German Cancer Research Center (DKFZ), Heidelberg, Germany; Department of Clinical Pathology, Suez Canal University, Ismailia, Egypt; Department of Radiation Oncology, University Medical Center Göttingen, Göttingen, Germany

**Keywords:** carbon ion irradiation, CTLA4, immunotherapy, PD-L1, radiotherapy, radioimmunotherapy, EO771 tumor

## Abstract

Radiotherapy can act as an *in situ* vaccine thereby activating tumor-specific immune responses that prevent tumor outgrowth in treated patients. While carbon ion radiotherapy has shown superior biophysical properties over conventional photon irradiation, the immunological effects induced have remained largely uncovered. The combination of radiotherapy with immune checkpoint inhibition (radioimmunotherapy) aims at further enhancement of anti-tumor immunity; however, studies on the immune cell composition in irradiated and distant tumors following radioimmunotherapy with carbon ions are scarce. We have established a bilateral tumor model by time shifted transplantation of murine, Her2+ EO771 tumor cells onto the flanks of immune competent mice followed by selective irradiation of the primal tumor, while sparing the consecutive tumor. We demonstrate that αCTLA4-but not αPD-L1-based radioimmunotherapy induces complete tumor rejection in our model. Intriguingly, local tumor control caused *in situ* immunization resulting even in eradication of non-irradiated, distant tumors. Moreover, cured mice were protected against EO771 rechallenge indicative of long lasting, tumor-protective immunological memory. Deconvolution of the treatment induced immunological effects by single cell RNA-sequencing (scRNA-seq) and concomitant flow cytometric analyses revealed in irradiated tumors predominating myeloid cells that developed into distinct tumor-associated macrophage clusters with upregulated expression of TNF and IL1 responsive genes, as well as activation of NK cells. Non-irradiated tumors showed higher frequencies of naïve T cells in irradiated mice, which were activated when combined with CTLA4 blockade. In conclusion, radioimmunotherapy with carbon ions plus CTLA4 inhibition reshapes the tumor-infiltrating immune cell composition and can induce complete rejection even of non-irradiated tumors. Our data present a rationale to combine radiotherapy approach with CTLA4 blockade to achieve durable anti-tumor immunity. Evaluation of future radioimmunotherapy approaches should thus not only focus on the immunological impacts at the site of irradiation but should also consider systemic immunological effects that might affect outgrowth of non-irradiated tumors.

## Introduction

Activating the patient’s autologous immune system to fight tumors has revolutionized the management of cancer (Cappell *et al*, 2020; Hodi *et al*, 2010; Larkin *et al*, 2019). While a plethora of new immunotherapeutic approaches have been investigated in recent years, also standard therapies, which are safe and well established, have shown promising immunomodulatory potential (Fucikova *et al*, 2020; Vanmeerbeek *et al*, 2020). Radiotherapy, one such example, not only induces DNA damage and cell death in irradiated tumor cells, but may also initiate immunogenic cell death characterized by the release of tumor antigens and damage-associated molecular patterns (DAMPs) like type I IFNs, ATP, and HMGB1 (Galluzzi *et al*, 2020; Golden *et al*, 2014; Vaes *et al*, 2021) possibly priming an anti-tumor immune response (Deng *et al*, 2014; Krombach *et al*, 2019). Moreover, the upregulated expression of MHC molecules (Reits *et al*, 2006; Schroter *et al*, 2020) and adhesion molecules (Rodriguez-Ruiz *et al*, 2017) as well as the release of immunostimulatory cytokines (Lugade *et al*, 2008) and chemokines (Matsumura & Demaria, 2010) create a pro-inflammatory tumor microenvironment (TME).

Remarkably, radiation-induced immune cell activation is not restricted to locally irradiated tumors but may also stimulate systemic immune responses that can affect distant metastasis (so called “abscopal effect”). Although the first radiation-induced abscopal effect was already described in the 1950s (Mole, 1953), only few cases have been reported to date (Abuodeh *et al*, 2016). This highlights the great impact of the immunosuppressive TME, which can be even further enhanced by radiotherapy (RT) inducing the expression of immune checkpoint molecules (Dovedi *et al*, 2014; Wirsdorfer *et al*, 2016) and the accumulation of suppressive immune cells like Tregs (Qu *et al*, 2010) as well as M2-like tumor-associated macrophages (TAMs) (Stafford *et al*, 2016). To counteract immunosuppression and promote systemic tumor immunity, preclinical and clinical studies meanwhile focus on radioimmunotherapy approaches combining RT with administration of immune checkpoint inhibitors against PD-1/PD-L1 and CTLA4 (Dewan *et al*, 2009; Dovedi *et al*., 2014; Golden *et al*, 2015; Hiniker *et al*, 2016; Kang *et al*, 2016; Rückert *et al*, 2021).

While immunomodulatory and therapeutic effects of radioimmunotherapy are well established for conventional photon RT with x-rays or γ-rays, little is known when it comes to particle radiation using protons or carbon ions. Notably, carbon ion radiation holds superior physical and biological characteristics over photon radiation (Chiblak *et al*, 2019; Marcus *et al*, 2021). Thus, the maximal carbon ion dose is released at the end of the beam track, called Bragg peak, allowing focused energy release. This enables precise irradiation of tumors located at deeper areas within the body, while sparing healthy tissue (Schulz-Ertner & Tsujii, 2007). Moreover, on the cellular level carbon ion irradiation induces a higher degree of complex irreversible DNA damage (Hagiwara *et al*, 2017; Lopez Perez *et al*, 2019), which is less dependent on oxygen supply and cell cycle phase (Jiang, 2012; Nickoloff, 2015).

Here, we use the murine Her2^+^ breast cancer cell line EO771 (Le Naour *et al*, 2020) and directly compared the anti-tumoral effects induced by carbon ion RT and conventional photon RT. We applied biologically equivalent doses (GyRBE) based on previously reported *in-vitro* clonogenic survival of EO771 tumor cell line (Hartmann *et al*, 2020). Irradiation was combined with checkpoint inhibition by antibodies against CTLA4 and PD-L1. Moreover, we present for the first time comprehensive single cell RNA sequencing analyses on individual immune cell subsets in carbon ion irradiated tumors compared to immune cell infiltrates in non-irradiated, distant tumors located outside the irradiation field.

Our results strongly suggest that the development of successful radioimmunotherapy approaches will require critical evaluation of therapy-induced effects not only in irradiated tumors, but importantly also in distant tumors outside the irradiation field. This applies in particular to metastatic lesions that represent drivers of tumor development.

## Results

### Checkpoint blockade improves growth control of locally irradiated and distant tumors

We first investigated the effects of carbon ion (C12) radiotherapy plus immune checkpoint blockade on outgrowth of irradiated in-field (iF) tumors and distant, non-irradiated out-of-field (oF) tumors as outlined in Fig. 1A. For this, mice were bilaterally challenged by time-shifted injections with equal tumor cell numbers. When tumors had established, mice were stratified into four treatment groups based on the volume of iF and oF tumors (see Methods). Next day, iF tumors were irradiated with three fractions of 3.08 Gray (Gy) carbon ions, while distant oF tumors were spared. This irradiation dose was selected based on our previous *in-vitro* clonogenic survival data: 3.08 Gy carbon ions showed equivalent reduction of the survival fraction compared to 5 Gy photons (Hartmann *et al*., 2020). This corresponds to a relative biological effectiveness (RBE ∼ 1.62) for the endpoint clonogenic survival for carbon vs. photon.

**Fig. 1.**
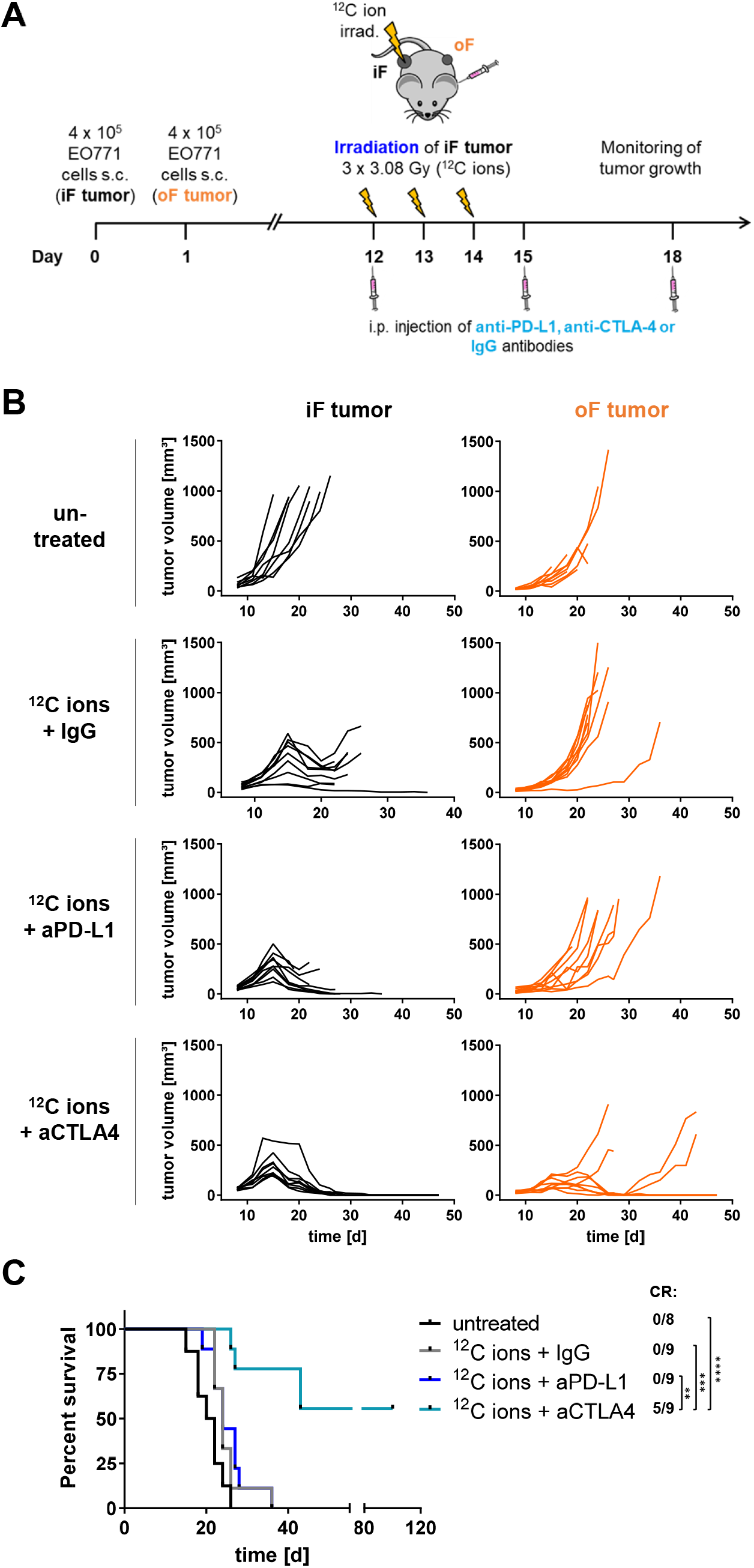
Tumor growth control after carbon ion RT combined with immune checkpoint blockade against PD-L1 or CTLA4. **(A)** Treatment schedule: C57BL/6 mice were bilaterally challenged by time-shifted injections (s.c.) with 4 × 10^5^ EO771 cells. When tumors had established, mice were randomized into treatment groups based on the volume of in-field (iF) and out-of-field (oF) tumors. Then, the iF tumor was irradiated with 3.08 Gy carbon ions on three consecutive days. On day 12, 15, and 18, mice received i.p. injections of blocking antibodies against PD-L1 or CTLA4. Control mice were injected with IgG isotype controls. **(B)** Individual growth curves of iF and oF tumors of untreated control mice or mice treated as indicated. **(C)** Kaplan-Meier survival analysis. In addition, complete response (CR) rates for each treatment group are shown. Significance was determined using a log-rank (Mantel-Cox) test with Holm-Bonferroni correction.

Moreover, mice received three injections with blocking antibodies against the checkpoint molecules PD-L1 or CTLA4, three days apart starting on the day of the first carbon ion irradiation. Following this treatment schedule, monotherapy with carbon ion irradiation resulted in growth delay and regression of iF tumors, while outgrowth of oF tumors appeared largely unaffected. Eventually, the majority of iF tumors grew out after a transient lag phase, however, further progression of these tumors could not be monitored due to the rapid outgrowth of non-irradiated tumors. Radioimmunotherapy with carbon ions combined with PD-L1-blocking antibody could further improve the curative effect on iF tumors, but hardly affected the growth of oF tumors. In contrast, mice receiving radioimmunotherapy with carbon ions plus CTLA4-blocking antibody rejected half of the oF tumors in addition to complete remission of iF tumors (Fig. 1B). Notably, this treatment regimen caused durable complete responses and improved survival in five out of nine mice (Fig. 1C). Thus, our bilateral tumor model suggests that CTLA4-blocking therapy is superior to PD-L1-blocking when combined with radiotherapy.

### Radioimmunotherapy with carbon ions or photons causes similar tumor control

Carbon ion irradiation is characterized by favorable physical and biological properties compared to conventional photon irradiation (Schulz-Ertner & Tsujii, 2007). Consequently, the therapeutic effects of both radiation types combined with CTLA4-blocking antibody were compared in the bilateral EO771 tumor model. For this purpose, BEDs of 5 Gy photons and 3.08 Gy carbon ions were applied, respectively (Hartmann *et al*., 2020), together with CTLA4 blockade as indicated in Fig. 1A.

Radioimmunotherapy with CTLA4-blocking antibody resulted in efficient tumor growth control leading to complete regression of 60% and 40% of both tumors with C12 and photon irradiation, respectively (Fig. 2A). Induced regressions were durable, lasting for more than 80 days resulting in a significant survival benefit compared to untreated mice or to mice treated with RT alone (Fig. 2B, C). Monotherapy with photons appeared slightly more efficient in controlling outgrowth of iF tumors compared to carbon ion RT (Fig. 2A). Yet, no therapeutic effect on the outgrowth of distant oF tumors was observed, irrespective of the irradiation type applied. Consequently, the survival rate of mice was comparable after monotherapy with photon and carbon ion RT, respectively, but was significantly improved when compared to untreated mice due to growth delay of iF tumors (Fig. 2B, C).

**Fig. 2.**
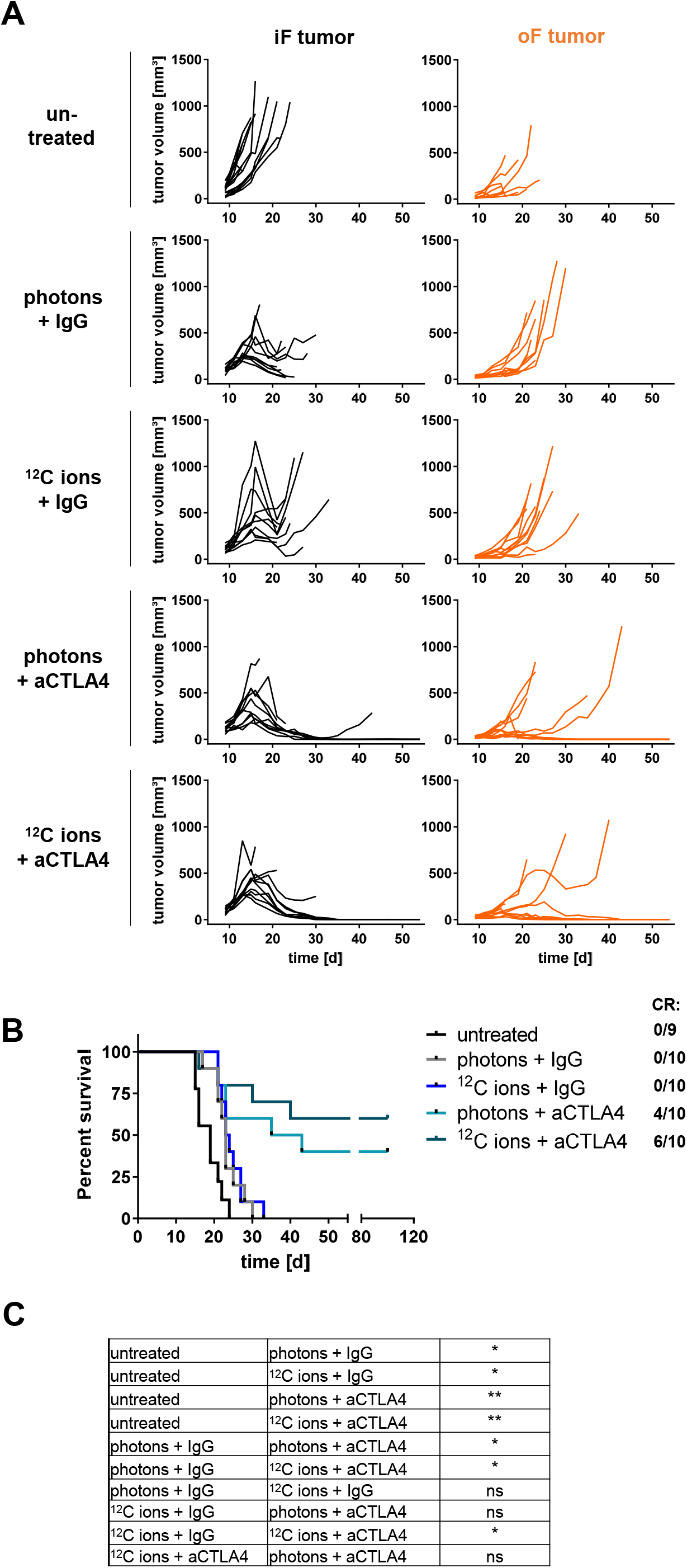
Irradiation with carbon ions or photons exhibits equal therapeutic capacity. **(A)** Individual growth curves of in-field (iF) and out-of-field (oF) tumors of untreated control mice or mice treated as indicated. **(B)** Kaplan-Meier survival analysis showing complete response rates for each treatment group. **(C)** Comparison of groups depicted in the Kaplan-Meier survival analysis. Significance was determined using a log-rank (Mantel Cox) test with Holm-Bonferroni correction.

Following radioimmunotherapy with photons and anti-PD-L1 antibody, results were comparable to those obtained in the carbon ion RT experiment (Fig. S1A, B and Fig. 1). Thus, addition of anti-PD-L1 antibodies to photon RT slightly improved growth control of iF tumors compared to photon RT alone, while the effect on oF tumors was neglectable. Monotherapy with immune checkpoint-blocking antibodies was largely inefficient in the bilateral EO771 model (Fig. S1A, B; Fig. S2A, B). In summary, combination of RT with CTLA4-blocking antibody was superior to the combination therapy with PD-L1 blockade and resulted in long lasting growth delay of both iF and oF tumors. Moreover, mice receiving carbon ion-based RT combined with anti-CTLA4 treatment showed best survival benefit (Fig. 1C, Fig. 2B).

### Composition of tumor-infiltrating immune cells following carbon ion-based radioimmunotherapy

Having observed durable therapeutic effects of carbon ion irradiation plus CTLA4-blocking antibody treatment on the survival of tumor bearing mice, we investigated the immune cell infiltrates within iF and oF tumors following carbon ion-based radioimmunotherapy in closer detail. Thus, mice were treated as outlined in Fig. 3A and tumors were harvested at day 18 to obtain animals with similar tumor sizes in all groups (Fig. 3C). Dissociated tumors from each treatment group were split into 2 halves: one half underwent flow cytometric analysis on CD45 gated immune cells for each individual tumor. The other half was pooled, sorted for CD45 positive cells, and subjected to single cell RNA sequencing (scRNA-seq) (Fig. 3A). The comprehensive sequencing dataset thus included 37441 tumor infiltrating immune cells. A first clustering resulted in 13 clusters with similar immune cell subpopulations as described before in other murine tumor models (Gubin *et al*, 2018; Zhang *et al*, 2020). Based on published gene expression profiles, these clusters were attributed to two main immune cell populations, i.e., *Itgam*-expressing TAMs (Xu *et al*, 2021) comprising TAM-A and TAM-B subtypes (Fig. 3B, D, E), and *Cd3e*-expressing T cells (Fig. 3B, E). Two further prominent populations consisting of *Cd19-*expressing B cells (Fig. 3B, D, E) and of neutrophils showing expression of *Csf3r (Zhang et al*., *2020)* were determined (Fig. 3B, E). Expression analysis of subtype associated marker genes enabled further differentiation of these immune cell sub-populations (Fig. S3A).

**Fig. 3.**
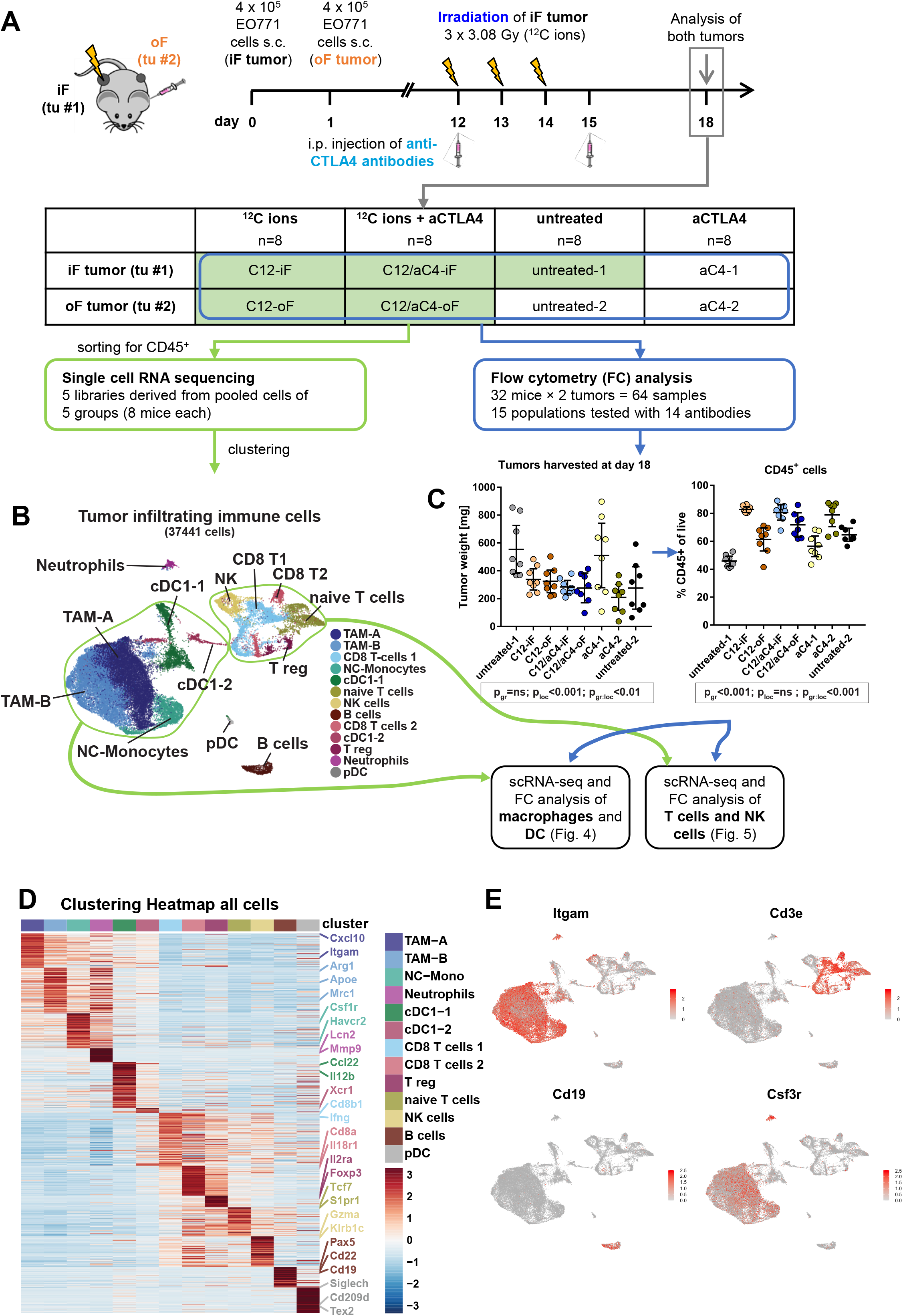
Characterization of the tumor-infiltrating immune cell composition following carbon ion-based radioimmunotherapy. **(A)** treatment schedule **(B)** UMAP embedding of 37441 tumor infiltrating cells of all conditions identifies immune cell subtypes; expression of marker genes is reported in Fig. 3 S1A. **(C)** FACS analysis of tumor infiltrating immune cells following treatment schedule as depicted in (**A**). **(D)** Differential expression heatmap of tumor infiltrating immune cells. Differential expression analysis revealed cluster specific genes that were ordered along clusters (Materials and Methods). Individual genes were annotated, according to the color code applied in Fig. (**B**). The table of the full heatmap is given in **Table S2**. **(E)** Tumor infiltrating immune cells include Macrophages/Monocytes (Itgam), T cells (Cd3e), B cells (Cd19) and Neutrophils (Csf3r). Statistics calculated using linear mixed models accounting for treatment groups and tumor locations (see Materials and Methods). p-values refer to type III analysis of treatment group (p_gr_), location (iF versus oF tumor, p_loc_), and their intercept (p_gr:loc_); complete statistics are reported in Table S10.

We noted that TAM-B cells outnumbered the other groups of immune cell populations, which was most pronounced in irradiated in-field tumors irrespective of CTLA4-specific antibody administration (Fig. S3B).

Of note, flow cytometry analysis revealed that iF tumors showed higher frequencies of CD45+ immune cells compared to oF tumors, irrespective of anti-CTLA4 treatment applied (Fig. 3C, right). To exclude potential effects of different tumor sizes we investigated the weights of the harvested tumors and found that they were comparable among irradiated mice both in iF and oF tumors (Fig. 3C, left). Thus, all treatment conditions seem to be at a similar disease stage allowing in-depth comparison of different immune cell subsets.

### Carbon ion irradiation reshapes the myeloid compartment in EO771 tumors

TAMs have been shown to interfere with local anti-tumor immune responses as reviewed in (Mantovani *et al*, 2017). Thus, we started to investigate the impact of carbon ion irradiation, with or without anti-CTLA4 administration, on gene expression in the intra-tumoral myeloid compartment. Sub-cluster analyses performed on *Itgam*-expressing immune cells revealed a continuum of myeloid cell types (Fig. 4A, B) similarly as described before (Gubin *et al*., 2018; Zhang *et al*., 2020) This continuum included *Chil3-*expressing monocytes (Fig. 4D), tissue-resident macrophages (TRMs) showing *Fcer1g* expression *(Dang et al, 2020)* (Fig. 4B), and a range of tumor associated macrophages designated as TAM-1 to TAM-7 (Fig. 4 A-C, F, Fig. S4). Additionally, we determined three dendritic cell (DC) clusters based on the expression of DC associated genes, namely cDC1-1 expressing *Ccl22*, cDC1-2 showing *Clec9a* expression, and *CD209a* expressing cDC2 cells (Zhang *et al*., 2020) (Fig. 4B, Fig. S4).

**Fig. 4.**
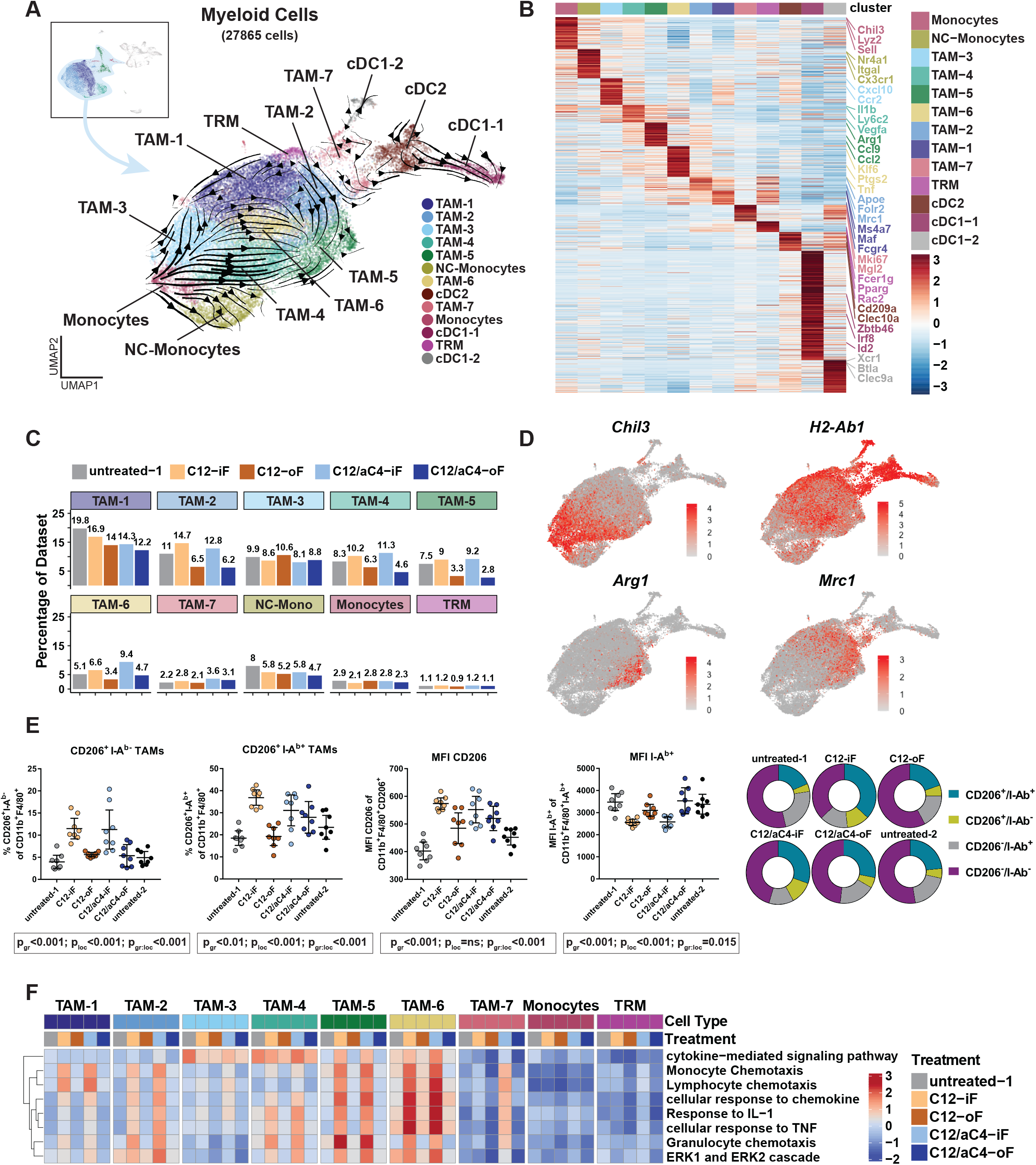
Direct carbon ion radiotherapy reshapes the TAM landscape. **(A)** Subclustering of 27865 myeloid cells of all treatment groups. Cells are embedded in UMAP space and RNA velocity estimation is plotted superimposed on top (black arrows). Two origins of differentiation are determined in TRM and Monocytes. Both give rise to trajectory endpoints in TAM-5 and between TAM-1 and TAM-3. Notably, monocytes additionally give rise to NC-monocytes. Besides Macrophages/Monocytes three clusters of classical DCs are found (UMAP of marker genes are shown in Fig. S4). **(B)** Differential expression heatmap of myeloid cell subclustering. Differential expression analysis revealed cluster specific genes that were ordered along clusters (see Materials and Methods). Individual genes were annotated, according the color code applied in Fig. 4A. The table of the full heatmap is reported in **Table S3**. **(C)** Cluster contribution of treatment groups to macrophage and monocyte clusters. iF clusters (C12-iF and C12/aC4-iF) show enrichment of clusters along the lineage towards TAM-5 (TAM-2, TAM-4, TAM-6, TAM-5, see Fig. S5A for aggregated percentages), while the origins of differentiation (Monocytes, TRM) as well as the other lineages (NC-Mono and towards TAM-1 and TAM-3) were unchanged. Percentages of DCs are shown in Fig. S5B. **(D)** Selected genes in myeloid cell subclustering. *Chil3* was expressed in cells from the monocyte lineage. *H2-Ab1* (=I-Ab) is enriched on the TAM-1 differentiation, while the TAM-5 lineage is enriched in *Arg1* and *Mrc1* (=CD206) TAMs. **(E)** Flow cytometric analysis of the myeloid immune cell compartment in tumors following the treatment schedule depicted in Fig 3A. Statistics calculated using linear mixed models accounting for treatment groups and tumor locations (see Materials and Methods). p-values refer to type III analysis of treatment group (p_gr_), location (iF versus oF tumor, p_loc_), and their intercept (p_gr:loc_); complete statistics are reported in Table S10. **(F)** Expression heatmap of enriched GO-terms in Macrophage/Monocyte clusters. Gene set enrichment analysis (GSEA) was used to infer enriched GO-terms from differential gene expression analysis (Fig. S6A and B). Selected terms were aggregated per cell type and treatment condition. iF conditions (C12-iF and C12/aC4-iF) show induction of the enriched terms especially in TAM-5 and -6. Other TAM groups show less induction while origins of differentiation (Monocytes and TRM) are unchanged. Heatmap of all clusters including dendritic cells is reported in Fig. S6C and D.

Next, we performed RNA velocity analysis to investigate the functional dynamics underlying the continuum of TAM subtypes determined. Focusing on macrophage and monocyte clusters we noted two opposite developmental origins represented by TRMs (top cluster) and by monocytes (bottom left cluster), respectively (Fig. 4A). Notably, developmental trajectories from both origins culminated in the same opposing TAM populations, namely TAM-5 and in between TAM-1 and TAM-3. At the same time, only monocyte-derived trajectories gave rise to *Nr4a1* expressing non-classical monocytes (NC-Monocytes). These data suggest that TAMs in our model derive from both local TRMs and from infiltrating monocytes. Furthermore, our data indicate a convergent development towards defined TAM states, as both TAM trajectory end points share contributions from both origins.

We then analyzed the effect of irradiation on the immune cell type compositions along the trajectories and found that the proportions of TRMs and monocytes, representing the origins of the TAM trajectories, remained unaffected irrespective of the radioimmunotherapy regimen applied (Fig. 4C). In contrast, carbon irradiated tumors (C12-iF and C12/aC4-iF) showed an increase of TAM-5 and of all TAM clusters along the trajectories approaching it (Fig. 4A, C). This included TAM-2 from TRM as well as TAM-4 and TAM-6 from monocytes (Fig. 4C, Fig. S5A). Notably, other TAM groups (TAM-1, TAM-3, TAM-7), non-classical monocytes (NC-Monocytes), or dendritic cells (cDC1-1, cDC1-2, cDC2) were not increased in iF tumors (Fig. 4C, Fig. S5B, respectively).

The macrophage subtype composition induced by iF carbon ion RT was then compared to respective TAM clusters described in murine tumor models following immune checkpoint therapy or myeloid targeted treatment (Gubin *et al*., 2018; Zhang *et al*., 2020). Therefore, we constructed modules of co-expressed genes (Trapnell *et al*, 2014) on myeloid cells and correlated these modules to TAM datasets presented by Gubin et al. (Gubin *et al*., 2018) and Zhang et al. (Zhang *et al*., 2020), respectively (see Methods). These analyses suggested a continuum of TRM-and monocyte-derived subtypes observed in our results concordant with both reference datasets (Fig. S5C). In fact, the TAM populations increased in carbon ion irradiated iF tumors, namely TAM-5 together with the more immature clusters TAM-4, and TAM-6 showed high similarity to previously published macrophages Mac_s5 and mmM15 involved in tumor rejection upon combined immune checkpoint therapy (Gubin *et al*., 2018; Zhang *et al*., 2020) (Fig. S5C). Furthermore, the developmental endpoint that was increased after C12 radiotherapy (TAM-5) showed expression of *Folr2*, a gene recently found to be expressed in TRM-derived anti-tumor TAMs (Nalio Ramos *et al*, 2022) (Fig. S5D).

The observed changes among the myeloid compartment in tumors following direct carbon ion irradiation were validated by flow cytometric analyses performed on TAMs from individual tumors (Fig. 4E). This analysis revealed increased proportions of CD206 (*Mrc1*) expressing TAMs following ∼ 3.1 Gy carbon ion irradiation, independent of anti-CTLA4 treatment, while the surface expression levels of IA^b^ (*H2-Ab1*) were reduced in iF tumors (Fig. 4E). Of note, a similar pattern was observed when iF tumors were irradiated with 5 Gy photons (Fig. S5E and F).

Thus, our data suggest that carbon ion-based radiotherapy induces specific changes in the cellular composition of myeloid cells within the continuum of TAM differentiation.

### Carbon ion-based radioimmunotherapy shapes TAM responses

CD206-expressing TAMs have been generally considered as anti-inflammatory macrophage subtype promoting tumor growth (Mantovani *et al*., 2017). However, recently TAMs expressing both CD206 and FOLR2 were shown to be involved in breast cancer rejection (Nalio Ramos *et al*., 2022). We noticed increased frequencies of CD206-expressing TAMs in iF tumors whose outgrowth was prevented upon carbon ion RT (Fig. 1, Fig. 2, Fig. 4E). Moreover, CD206 surface expression was most pronounced on TAMs from iF tumors, whereas IA^b^ expression appeared significantly reduced (Fig. 4E). We therefore hypothesized that besides an increase in specific clusters, iF carbon ion RT might alter gene expression within TAMs. Hence, we performed differential gene expression analysis of myeloid cells comparing all in-field (data from C12-iF and C12/aC4-iF) against untreated tumors and found strong induction of cytokine and chemokine expression, including *Ccl2, Ccl7, Cxcl2* and *Cxcl3* among myeloid cells of iF tumors (Fig. S6A). Gene set enrichment analysis (GSEA) revealed a significant enrichment of chemotaxis and migration terms, as well as ERK signaling and cellular responses to TNF-alpha and IL-1 (Fig. S6B). To determine the cell types showing induction of the aforementioned pathways, we aggregated expression per cluster and treatment condition (Fig. 4F). This confirmed induction of chemotaxis, ERK signaling and responses to IL-1 as well as TNF-alpha of iF tumors compared with oF or untreated tumors, which was most pronounced in TAM-5 and TAM-6. Moreover, induction of the same gene expression pattern was consistently observed at lower level also in TAM-1, TAM-2, and TAM-4 clusters of iF tumors. In TAM-7 clusters of iF tumors, induced expression of these genes appeared dependent on anti-CTLA4 antibody administration. Notably, the expression profiles in the developmental origins, TRM and monocytes, were not altered between the treatment groups.

The modified gene expression patterns observed were restricted to macrophages, since neither dendritic cells, nor NK cells or T cells showed induction of the GSEA terms observed among the TAMs (Fig. S6C and D). Of note, neutrophils, showed a similar pattern as macrophages (Fig. S6D), however this population represented less than 2% of the total immune cell infiltrate (Fig. S3B). We thus propose that the gene expression profiles determined in iF tumors following carbon ion irradiation was restricted to TAM subsets.

To validate whether changes in gene expression of TAMs from iF tumors were associated with carbon ion irradiation therapy or were affected by other treatments, we aggregated all macrophages per treatment condition and compared them to TAMs derived from tumors after immune checkpoint therapy i.e., without radiotherapy applied (Gubin *et al*., 2018). Most interestingly, only iF irradiation, but not immune checkpoint therapy alone induced pathways affecting response to IL-1, TNF-alpha and chemokines as well as, chemotaxis and ERK signaling (Fig. S6E). Combination therapy even increased these effects (compare C12-iF vs. C12/aC4-iF TAMs; Fig. S6E).

Finally, to test whether anti-CTLA4 treatment induced gene expression changes irrespective of carbon ion irradiation, we compared C12 (iF and oF) with C12/aC4 (iF and oF) TAMs. Differential gene expression analysis revealed only two genes increased in C12/aC4 including *Cxcl9* (Fig. S6F), which has been recently described as chemokine preferentially expressed by TAMs of triple negative breast cancer patients responding to immune checkpoint blockade therapy (Zhang *et al*., 2020).

Thus, our data suggest that carbon ion-based radiotherapy induces changes in both cell composition and gene expression patterns specifically among TAMs. Combination with anti-CTLA4 antibody treatment further potentiates these effects.

### Carbon ion-based radioimmunotherapy activates CD8^+^ T cells in non-irradiated tumors

The T cell / NK cell compartment represented another dynamic population of tumor-infiltrating immune cells (Fig. 3B, Fig. S3A, B). Dimensionality reduction in UMAP space revealed ten clusters including naïve CD4^+^ and CD8^+^ T cells (*Cd4 or Cd8, S1pr1, Il6ra, Il7r* and *Lef1*), Th1 cells (*Tnfsf8* and *Tnfsf11*) and regulatory T cells (Treg; *Ctla4, Tnfrsf4, Foxp3, Itgb8*) as well as a cluster of NK cells (*Gzma* and *Klrb1c*) (Fig. 5A, C, Fig. S7). Additionally, we found different clusters of effector T cells (Fig. 5A), namely dividing effector cells (Div. Eff.; *Mki67*) and dividing *Il18r1*-expressing CD8^+^ T cells (Div. IL18R CD8; *Mki67*), activated/exhausted CD8^+^ T cells (CD8 Act/Exh; *Bhlhe40, Pdcd1, Litaf*), CD8^+^ T effector memory cells (CD8^+^ EM; *Havcr2, Lag3, Cd160, Ifng, Cxcr6*), and CD8^+^ T cells additionally expressing *Il18r1* and *Il18rap* (Timperi *et al*, 2017) (Fig. 5B, C and Fig. S7). Of note, comparison of our clusters to a recently published reference T cell atlas (Andreatta *et al*, 2021) confirmed the different T cell subsets determined in our study (Fig. S8A).

**Fig. 5.**
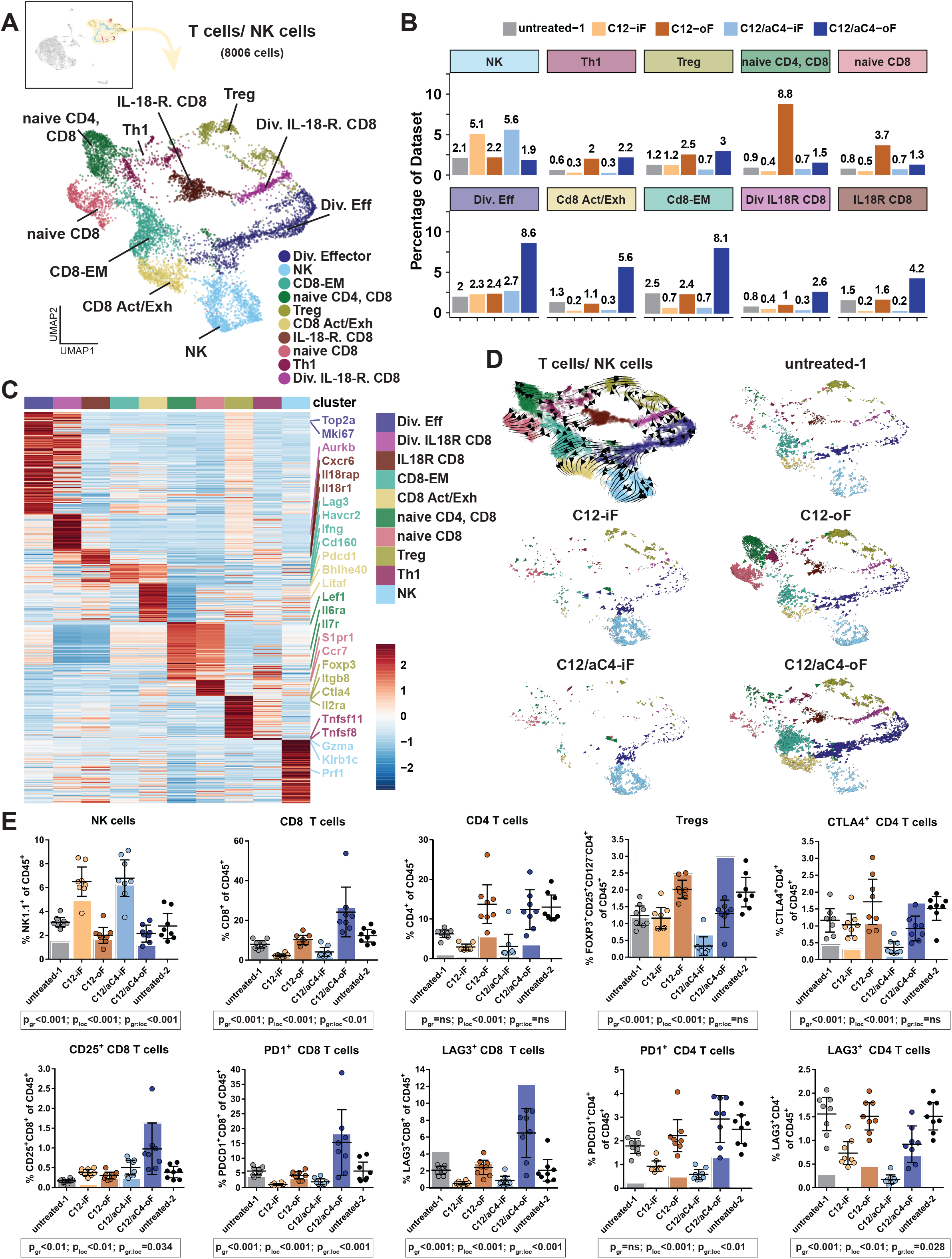
Radioimmunotherapy exerts opposed effects on NK cells and T cells in iF and oF tumors. **(A)** UMAP embedding of subclustering of lymphoid cells including 8006 T and NK cells from all treatment conditions. UMAP with marker genes are depicted in Fig. S7. **(B)** Cluster contribution of different treatment conditions to T and NK cell clusters reveals increase of NK cells in iF conditions (C12-iF and C12/aC4-iF). C12-oF tumors are enriched for naïve T cells (naïve CD4 and CD8), while C12/aC4-oF show enrichment for activated and proliferating cell types (bottom row). **(C)** Differential expression heatmap of lymphoid cell subclustering. Differential expression analysis revealed cluster specific genes that were ordered along clusters (Materials and Methods). Individual genes were annotated, following the color code of Fig. 5A. The table of the full heatmap is reported in **Table S7**. **(D)** RNA velocity embedded in UMAP space for lymphoid cells. Left top shows all conditions in UMAP space with velocity arrows superimposed in black. Other panels show individual treatment conditions with each cell depicted as an arrow with size and direction relative to velocity. Note that iF conditions (C12-iF and C12/aC4-iF) show velocity from dividing cells towards NK cells. The C12-oF condition shows some velocity towards activated CD8^+^ cells, while most cells remain naïve. In contrast, C12/aC4-oF shows strong velocity from dividing cells towards activated cells with few naïve T cells or NK cells. **(E)** Proportion of tumor infiltrating immune cell subpopulations determined by flow cytometric analysis in tumors from individual mice (dots) and by scRNA sequencing of pooled tumors (columns), following the treatment schedule depicted in Fig 3A. Statistics calculated using linear mixed models accounting for treatment groups and tumor locations (see Materials and Methods). p-values refer to type III analysis of treatment group (p_gr_), location (iF versus oF tumor, p_loc_), and their intercept (p_gr:loc_); complete statistics are reported in Table S10.

We next investigated the effects of iF vs oF irradiation on the intratumoral T cell and NK cell composition using scRNA data and flow cytometric analysis. In fact, iF tumors showed an increase of NK cells upon carbon ion irradiation and carbon ion-based radioimmunotherapy (C12-iF and C12/aC4-iF groups) compared to untreated and oF tumors (Fig. 5B, E, Fig. S8B).

On the other hand, depending on co-administration of anti-CTLA4 antibody, oF tumors were increased in naïve (C12-oF) or activated CD8^+^ T cells (C12/C4-oF), respectively. Thus, combination of carbon ion irradiation and anti-CTLA4 treatment massively enhanced the activated effector cell clusters as seen for dividing effector cells, CD8^+^ activated/exhausted T cells, CD8^+^ T_EM_ cells, dividing and non-dividing CD8^+^ IL-18R^+^ T cells (Fig. 5B). Notably, dividing effector cells showed expression of CD8^+^ T-cell markers (*Cd8a*) in C12-oF and C12/aC4-oF tumors, while in iF tumors the dominant immune cell population was represented by NK cells expressing *Gzma* and *Klrb1c* (Fig. 5D, S7, S8D). Furthermore, we found an enhanced proportion of Treg cells in oF tumors (Fig. 5B, D), however due to a general increase of T cells, the relative abundance of Treg remained constant among the treatment groups (Fig. S8C). Similarly, irradiation increased the relative abundance of Treg in iF tumors, which was completely compensated by the combination treatment. Next, RNA velocity analysis was applied to investigate radioimmunotherapy-induced changes in developmental trajectories in the T cell / NK cell compartment. Embedding of velocities in UMAP space revealed that within iF tumors (C12-iF and C12/aC4-iF), the dividing effector cell cluster developed towards NK cells, but not towards activated CD8^+^ T effector cell (Fig. 5D). In C12-oF tumors the majority of T cells remained naïve, while a small group of cells showed a trajectory towards activated CD8^+^ T cell populations (Fig. 5D). In striking contrast, following radioimmunotherapy, T cells in C12/aC4-oF tumors showed strong velocities towards activated CD8^+^ T effector cell populations, while both naïve T cells and NK cells were rare (Fig. 5D). Taken together, we propose that combination therapy of carbon ion irradiation and anti-CTLA4 administration results in proliferation and activation of CD8^+^ T cells in distant oF tumors.

### Radioimmunotherapy activates in-field NK and out-of-field T cells

We then validated changes in the composition of the T cell / NK cell compartment determined through scRNA-seq by comparative flow cytometry analyses performed on infiltrating immune cell populations of individual tumors. Immunofluorescence staining against NK1.1 (encoded by *Klrb1c*) confirmed the increase of NK cells within iF tumors, irrespective of anti-CTLA4 treatment. Moreover, consistent with our scRNA-seq results shown in Fig. 5B, increased proportions of CD8^+^ T cells and activated CD8^+^ T cells expressing LAG3 or PD1 were only detected in oF tumors of mice that had received carbon ion radiation and anti-CTLA4 treatment, further supporting the activation of this T cells upon combination therapy (C12/aC4-oF, Fig. 5E). Of note, similar infiltration patterns of NK cells and T cells were seen within iF tumors following irradiation with photons, again irrespective of co-administered anti-CTLA4 antibody (Fig. S8E). However, the selective increase of exhausted and activated CD8^+^ T cells in oF tumors after radioimmunotherapy with photons (Ph/aC4-oF) was less pronounced compared to carbon ion-based radioimmunotherapy. Finally, we investigated CTLA4 expression in T cells. While scRNA-seq data revealed many cells expressing *Ctla4* on RNA level (Fig. S7), FACS analysis showed that the proportion of CD4^+^ T cells with CTLA4 surface protein expression was reduced upon anti-CTLA4 application in both iF and oF tumors (Fig. 5E). These data support that anti-CTLA4 antibody treatment efficiently depletes CTLA4 expressing Tregs (Simpson *et al*, 2013).

As carbon ion irradiation combined with anti-CTLA4 treatment increased activated CD8^+^ T effector cell populations within oF tumors (Fig. 5B, E), we investigated radioimmunotherapy-associated effects on gene expression among tumor infiltrating CD8^+^ T cells. While more abundant, activated CD8^+^ T cell populations did not differ in their gene expression patterns, when comparing C12-oF and C12/aC4-oF tumors (Fig. S9A and Supplementary Table S9). In contrast, naïve CD8^+^ T cells showed increased expression of a large set of genes following combined therapy with carbon ion irradiation and anti-CTLA4 treatment (Fig. S9B). GSEA revealed enrichment for terms such as lymphocyte differentiation and mononuclear cell differentiation as well as positive regulation of cytokine production (Fig. S9C). Comparing the expression of enriched terms and other T cell related GO-terms we found that upon C12/aC4 treatment, the naïve CD8^+^ T cell cluster (Fig. 5A, magenta), but not the naïve CD4^+^/CD8^+^ T cell cluster (Fig. 5A, green) showed increased expression of GO-terms related to T cell activation and differentiation (Fig. S9D). Thus, while there was no difference in gene expression among activated T cells, monotherapy with carbon ion irradiation resulted in naïve T cells, whereas radioimmunotherapy including anti-CTLA4 treatment induced expression of genes associated with T cell activation and differentiation.

Taken together, our analyses uncover the specific effects of carbon ion-based radio-monotherapy and of CTLA4 targeted radioimmunotherapy, respectively, on tumor resident T cells and NK cells. While NK cells were increased in iF tumors, independent of co-administered anti-CTLA4 antibody, oF tumors showed a larger contribution of T cells. However, only the combined radioimmunotherapy regimen resulted in a strong activation of CD8^+^ T cells with reduced CTLA4 surface expression within the oF tumor. This suggests a cooperative effect of these two treatment modalities in controlling both primary and distant tumor growth.

### Radioimmunotherapy induces protective long-term immunity against EO771 tumors

To estimate whether a potent and long-lasting anti-tumor response was induced we continued to investigate the surviving mice. Animals showing complete responses after photon or carbon ion RT combined with CTLA4 checkpoint blockade were re-challenged by subcutaneous injection of EO771 tumor cells three to six months after initial therapy. Strikingly, none of these mice showed tumor take, whereas five of six naïve mice developed tumors (Fig. 6A). Moreover, to test whether mice rejecting the secondary tumor had tumor-specific T cells we investigated splenocytes. Mice that had resisted secondary tumor challenge showed EO771-reactive T cell responses, which appeared more pronounced after radioimmunotherapy with photons compared to carbon ions, whereas no T cell responses against EO771 tumor cells were detectable in untreated control mice (Fig. 6B). Most importantly, the memory immune response was specific for EO771 tumor cells, as significantly lower amounts of INF-γ spots were detected when splenocytes were co-cultured with syngeneic cell lines of other tumor entities (Fig. 6C). Thus, taken together our data suggest that aside from generating a potent acute immune response of activated T cells in non-irradiated tumors, combination of carbon irradiation with anti-CTLA4 blockade produces long-lasting anti-tumor immunity sufficient to suppress future tumor outgrowth.

**Fig. 6.**
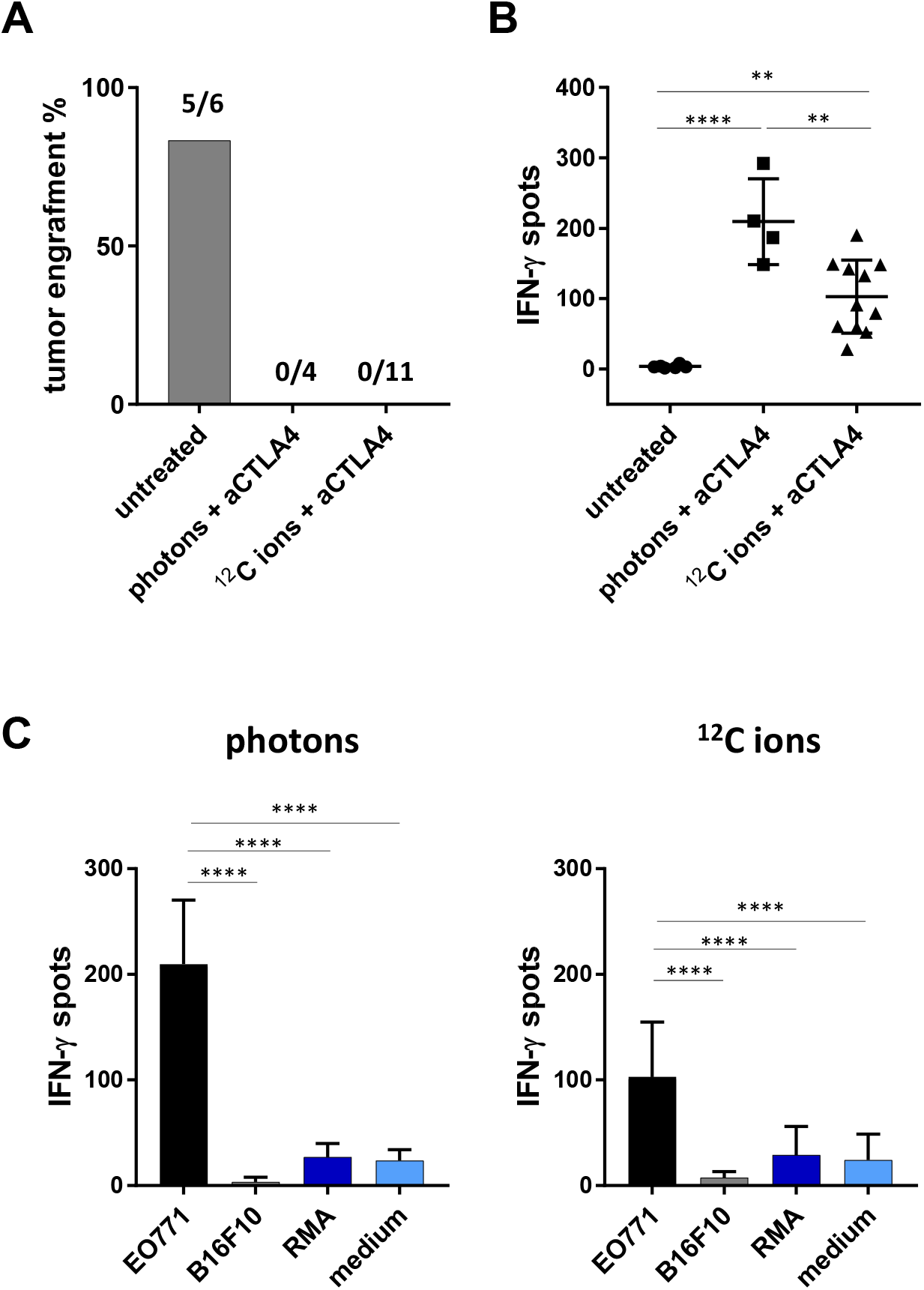
Mice cured by photon or carbon ion RT combined with CTLA4 checkpoint blockade show complete protection against EO771 tumor re-challenge. **(A)** Percentage of successful tumor engraftment in untreated mice or in mice whose EO771 tumors had regressed upon treatment with photon or carbon ion RT, respectively combined with CTLA4 blockade. Numbers of tumor rejecting mice per animals engrafted are depicted above the columns. **(B)** INF-γ ELISpot assay with splenocytes from cured mice and naïve control animals co-cultured o/n with EO771 target cells. **(C)** Specificity of the T cell response was assessed by IFN-γ ELISpot assays performed after co-culture of splenocytes with syngenic tumor cells lines B16F10 and RMA in comparison to EO771 cells. Significance was determined by one-way ANOVA with post-hoc Turkey’s test.

## Discussion

Irradiation can act as an *in situ* vaccine creating a pro-inflammatory TME and thus offering great potential for the initiation of anti-tumor immune responses (Goto, 2019). While the majority of preclinical and clinical studies have focused on conventional photon irradiation, the immunomodulatory potential of carbon ion irradiation is still under investigation. A recent study investigated the therapeutic effects of C12 irradiation combined with co-inhibition of PD-L1 and CTLA4 in the murine LM8 osteosarcoma model. The authors could demonstrate tumor growth control in the combination group (Takahashi et al, 2019), however, the effects of individual checkpoint inhibitor blockade in combination with C12 irradiation were not investigated and analysis of immune cell infiltrates was restricted to T cells. In our study, we demonstrate that mice bearing bilateral EO771 tumors show complete responses in about half of the cases upon three times ∼3.1 Gy physical dose of carbon ion irradiation when combined with anti-CTLA4 blockade. Additionally, we compared the therapeutic effects of carbon ion RT to conventional photon RT, as carbon ion RT has been reported to hold favorable physical properties (Schulz-Ertner & Tsujii, 2007). Our data suggests that both modalities can induce strong and comparable anti-tumor immunity at Gy RBE doses. Moreover, our scRNA-seq data show that the CD45^+^ immune cell infiltrates were dominated by *Itgam* expressing myeloid cells and a lymphoid compartment comprising mainly T cells and NK cells.

Selective analysis of myeloid cells revealed multiple TAM clusters, as well as several monocyte and DC clusters. Interestingly, RNA velocity analysis suggested the presence of Ly6C^+^ infiltrating monocytes and Ly6C^-^ TRMs, respectively as developmental origins of the TAM spectrum determined. In fact, Ly6C^+^ blood monocytes have the capacity to extravasate into tissues (Mildner *et al*, 2017), thereby possibly founding the developmental origin of Ly6C^+^ monocytes in our model. Results of our RNA velocity analyses suggested eventual development into macrophages of the TAM-5 cluster and a cell state in between clusters TAM-1 and TAM-3.

In irradiated tumors we found TAM-2, -4, -5, and -6 clusters to be increased in cell numbers. Furthermore, carbon ion therapy induced expression of genes belonging to chemokine signaling, monocyte and lymphocyte chemotaxis, as well as response to IL-1 and TNF..

Monocytes and the TAM-6 cluster showed enhanced expression of *Nr4a1*, a transcription factor preferentially expressed in Ly6C^-^ monocytes (Mildner *et al*., 2017) and non-classical monocytes. In fact, Nr4a1-dependent Ly6C^low^ expressing monocytes with inflammatory capacity (Brunet *et al*, 2016) or intravascular scavenger function have been described (Carlin *et al*, 2013).

The TAM-5 cluster together with the developmentally adjacent TAM-4 and TAM-6 clusters were increased after irradiation in iF tumors. Since these clusters show high similarity with myeloid clusters involved in tumor rejection following immune checkpoint therapy, i.e. treatment without radiotherapy (Gubin *et al*., 2018; Zhang *et al*., 2020), we conclude that TAM-4 to TAM-6 might be involved in an ongoing anti-tumoral myeloid immune response. This is in line with expression of *Folr2* that was recently shown to be specifically expressed in TRM-derived anti-tumor TAMs (Nalio Ramos *et al*., 2022).

Our gene expression analysis revealed enhanced expression of *Ccl2, Ccl7, Cxcl2* and *Cxcl3* in TAMs of carbon ion irradiated tumors. CCL2 and CCL7 were shown to recruit immunosuppressive T cells and myeloid cells (Korbecki *et al*, 2020). Remarkably, CCL2 was described to have both tumor promoting as well as anti-tumor function (Jin *et al*, 2021). Indeed, CCL2 can drive M2-like polarization of TAMs (Fei *et al*, 2021) characterized by scavenger function (Mantovani *et al*., 2017), and in fact, CD206 positive macrophages were shown to be involved in the clearance of apoptotic cells *in vivo* (Gordon & Pluddemann, 2018). CXCL2 has been reported to recruit Gr1^+^ neutrophils, which in turn promoted tumor growth through impaired T cell homing, aberrant angiogenesis, and resistance to checkpoint blockade (Faget *et al*, 2017). Notably, the chemokines mentioned here have mainly been reported as products released by tumor cells, although expression of CCL2 and CCL7 by TAMs has been found as well (Korbecki *et al*., 2020).

While these chemokines have been described to be tumor promoting largely by eventual suppression of effector T cells, we found T cells underrepresented in iF tumors suggesting a minor role of T cells and possibly a direct function of scavenging TAMs and/or attracted NK cells during the tumor rejection process. In line with this notion, we found a marked increase in NK cells in iF tumors independent of co-administered anti-CTLA4 antibody, whereas no such increase was observed in oF tumors. Interestingly, enhanced migration of NK cells upon photon irradiation has been reported before and was explained by radiation induced CXCL16 release from tumor cells and CXCR6 surface expression on NK cells (Yoon *et al*, 2016). NK cells can be activated by various chemokines including IL1 produced by DC or macrophages and are known to release mediators such as IFNγ and TNF in return (Abel *et al*, 2018; Cooper *et al*, 2004). Notably both, TNF and IL1 responsive genes were among the induced terms in TAMs from iF tumors independent of anti-CTLA4 treatment. In fact, NK cell mediated killing of the EO771 tumor cell has been demonstrated (Collin *et al*, 2017; Liu *et al*, 2021; Mayfosh *et al*, 2021) and NK cells were shown to be relevant in growth control of EO771 tumors (Tu *et al*, 2017). This supports our speculation that NK cells activated upon tumor entry might have contributed to the eradication of iF tumors.

Furthermore, we observed a reduced abundance of T effector cells in iF tumors, which might be due to cytotoxic irradiation effects, while radio-resistant Treg (Qu *et al*., 2010) and possibly a small fraction of resistant tumor resident T cells (Arina *et al*, 2019) remained. We hypothesize that tissue damage caused by irradiation and NK cell activity might result in release of tumor antigens that are taken up by scavenging M2-like TAMs (Mantovani *et al*., 2017) and / or by dendritic cells. These antigen-fed myeloid cells could migrate to draining lymph nodes where they encounter a less immunosuppressive environment allowing them to prime EO771-specific T cell responses. Activated T cells then reach out to distant sites to eradicate oF tumors through abscopal effector function (Pevzner *et al*, 2021), thereby possibly explaining the high proportion of activated and exhausted CTLs in oF tumors. This scenario might also explain why increased CD8+ T cells were activated in oF tumors only when irradiation was combined with anti-CTLA4 antibody administration.

We selected anti-CTLA4 antibody as checkpoint inhibitor in our study, since co-administration of CTLA4 blocking antibody induced superior anti-tumor effects compared to anti-PD-L1 antibody in our initial radioimmunotherapy experiments. CTLA4-blocking antibodies act during early phase of cellular immune responses facilitating efficient T cell priming while counteracting the suppressive function of CTLA4 expressing Treg (Buchbinder & Desai, 2016). CTLA4 blockade may efficiently synergize with RT in several ways: for example, RT triggers the release of tumor antigens and DAMPs (Galluzzi *et al*., 2020; Golden *et al*., 2014; Vaes *et al*., 2021) enabling priming of anti-tumor immune responses that may be further enhanced by CTLA4 blockade. Moreover, this study and others (Qu *et al*., 2010) showed that RT can induce a relative increase of Tregs, hence Treg depletion by anti-CTLA4 treatment may be of particular relevance. Co-application of PD-L1 blocking antibody with carbon ion RT showed anti-tumor effects on iF tumors, while outgrowth of oF tumors remained unaffected. Possibly, expression of PD-1 and PD-L1, respectively, by immune cells within the TME and / or by EO771 tumor cells was below threshold to enable effective anti-PD-L1 mediated rejection of oF tumors.

Studies investigating the composition and transcriptome of tumor infiltrating immune cells upon particle irradiation are scarce. One report describes immune cell infiltration and differential expression of 25 “immune response genes” upon monotherapy with 16.4 Gy protons in the murine transplantable CT26 colon cancer model (Mirjolet *et al*, 2021). More recently, the synergizing effect of a bifunctional protein simultaneously blocking TGFβ and PD-L1 combined with photon irradiation in reprogramming the immune suppressive TME and inducing abscopal effects was demonstrated in the murine 4T1 breast cancer model (Lan *et al*, 2021). This approach also prevented radiation induced lung fibrosis, thus scRNA-seq revealed reduced expression of fibrosis associated gene clusters in dissociated lung tissues of treated mice (Lan *et al*., 2021). However, the investigations mentioned above neither focused on carbon ion irradiation, nor did they include separate scRNA-seq analyses on sorted tumor infiltrating lymphocytes and TAM subpopulations, as presented in our study.

Irradiation, especially when combined with immunotherapy, acts not only directly on irradiated (iF) tumors, but also triggers indirect immunological responses that might result in abscopal effects (Pevzner *et al*., 2021). Notably, we could evaluate both direct (iF) and indirect (oF) effects of various radio-immunotherapy regimens on outgrowth of local and distant tumors, respectively, within the same animal. This advantage over studies merely focused on irradiated tumors, might provide new insights for the development of clinical radioimmunotherapy trials.

Finally, our re-challenge experiments showed that both photon and carbon ion RT with CTLA4 blockade induced a robust tumor-specific memory response. Thus, 100% of mice having shown complete responses against EO771 tumors after RT plus CTLA4 blockade rejected a secondary EO771 tumor engraftment and splenocytes of these mice showed EO771-specific INFγ responses. Taken together, our comprehensive investigation of the bilateral EO771 tumor model shows that carbon ion irradiation combined with anti-CTLA4 antibody administration has significant impacts on tumor-infiltrating immune cells: i) myeloid cells predominate in-field tumors and might develop into a distinct TAM cluster that is constantly replenished by TRMs and infiltrating monocytes; ii) in-field TAMs upregulate genes involved in chemotaxis and responses to TNFα and IL-1; iii) NK cells are activated in iF tumors; iv) whereas in oF tumors T cells are increased and activated, possibly via abscopal effects.

Furthermore, we demonstrate that CTLA4-based radioimmunotherapy with carbon ions or photons are effective in inducing rejection of both local and distant EO771 tumors, as well as in generating immunological memory against tumor re-challenge. Carbon ion irradiation has been used in around 30 clinical trials to date, primarily as monotherapy or in conjunction with chemotherapy (clinicaltrials.gov). Only two trials contain immune-checkpoint inhibition based on PD-1 blocking antibodies; however, the status of these trials is either uncertain or in the recruitment phase. Notably, the current study discovered the immunological advantages of combination treatment in out-of-field tumors, which is clinically relevant because non-irradiated metastases drive disease progression and mortality. Our findings should be validated in additional murine tumor types and confirmed in the human system before they can be translated. Although more research is needed, our results demonstrate that evaluating the radioimmunotherapeutic efficacy of treatment regimens necessitates consideration of both irradiated (iF) and distant (oF) tumors. Only by taking all these immunological effects into account can the full potential of a therapy regimen be determined.

## Material and Methods

### Cell lines and cell culture

The murine cell line EO771 was originally established from a spontaneous breast cancer of a C57BL/6 mouse and was recently characterized as estrogen receptor α negative, estrogen receptor β positive, progesterone receptor positive and ErbB2 (Her2) positive (Le Naour *et al*., 2020). EO771 cells were purchased from TEBU-Bio (Offenbach, Germany) and were cultured in RPMI 1640 medium (Thermo Fisher Scientific, Waltham, USA) supplemented with 10% heat-inactivated FCS, 10 mmol/L HEPES (Thermo Fisher Scientific), 100 U/ml Penicillin, and 100 μg/ml Streptomycin (Thermo Fisher Scientific). B16F10 cells (ATCC, Manassas, USA) and RMA cells (kindly provided by G. Hämmerling, DKFZ) were cultured in RPMI 1640 supplemented with 10% heat-inactivated FCS, 100 U/ml Penicillin, and 100 μg/ml Streptomycin.

### Tumor model and treatment

Animal experiments were approved by the District Government in Karlsruhe, Germany (approval ID 35-9185.81/G-209/18). Animal care was provided by the center for preclinical research at the DKFZ. Female C57BL/6J mice (Janvier Labs, Le Genest-Saint-Isle, France) were implanted s.c. with 4 × 10^5^ EO771 cells into the right hind leg and the left flank, respectively one day apart. Tumor diameters were measured with a caliper every 2-3 days and tumor volume was calculated as follows: V [mm^3^] = largest diameter [mm] x (smallest diameter [mm])^2^/2. When tumors were established, mice were assigned to treatment groups stratified by tumor volume of both tumors. Mean tumor volumes of leg tumors ranged between 120-200 mm^3^, those of tumors in the left flank between 40-60 mm^3^. Carbon ion RT of murine tumors was performed at the Heidelberg Ion-Beam Therapy Center (HIT) with a horizontal beamline using the raster scanning technique. Irradiation doses were delivered within a 20 mm wide extended spread out Bragg peak (dose average linear energy transfer (LET), 103 keV/μm) adjusted with a 30 mm wide acrylic absorber. Carbon ion RT was locally delivered in 3 fractions of 3.08 Gy to the leg tumors (in-field tumors), while the mice were anesthetized with isoflurane. Photon RT was performed using a Faxitron MultiRad225 irradiator (Faxitron Bioptics, LLC, Tucson, USA). 5 Gy photons were applied per fraction, which was determined as biologically equivalent to 3.08 Gy carbon ions previously. During photon irradiation, tumor-bearing mice were drugged with an antagonizing anesthesia. To this end, mice were anesthetized with a mixture of Medetomidin hydrochloride (0.25 mg/kg, Zoetis, Parsippany-Troy Hills, USA) and Midazolam (2.5 mg/kg, Ratiopharm, Ulm, Germany) diluted in 0.9% NaCl, which was antagonized with a mixture of Atipamzole hydrochloride (1.25 mg/kg, Zoetis, Parsippany-Troy Hills, USA) and Flumazenil (0.25 mg/ml, Fresenius Kabi, Bad Homburg, Germany) diluted in 0.9% NaCl. During irradiation of the in-field (iF) tumor, the body of the mouse including the out-of-field (oF) tumor at the flank were protected by a lead shield (dose rate: 5.57 Gy/min).

Starting with the first fraction of RT, mice were injected with 100 µg of anti-CTLA4 antibody, anti-PD-L1 antibody, polyclonal Syrian hamster IgG, or rat IgG2b isotype control (all from Bio X Cell, West Lebanon, USA). Mice received three injections of checkpoint blocking antibodies three days apart.

As termination criteria largest diameter >15 mm, tumor ulceration, signs of severe illness such as apathy, respiratory distress, or weight loss over 20% were defined. Whenever one of these criteria was met, mice were sacrificed by cervical dislocation or gradual CO_2_ exposition.

### Enzymatic treatment of tumors and preparation of single cell suspensions

Excised tumors were cut into small pieces following by incubation 0.5 mg/ml collagenase D (Hoffmann-La Roche, Basel, Switzerland), 10 µg/ml DNAse I (Sigma-Aldrich), 0.1 µg/ml TLCK (Sigma-Aldrich) and 10 mM HEPES solution (Sigma-Aldrich) diluted in HBSS for 1 h at 37 °C while shaking at 200 rpm. Digested tumors were passed through a 70 µm cell strainer and pelleted single cell suspensions were incubated with ACK lysis buffer (Thermo Fisher Scientific) to remove erythrocytes. Cells were washed and resuspended in FACS buffer (PBS + 3% FCS) for immunofluorescence staining and flow cytometric analysis.

### Flow Cytometry

Single cells suspensions were incubated a mixture containing 50 µg/ml rat anti-mouse CD16/CD32 (BD Biosciences, Franklin Lakes, USA), normal Syrian hamster serum (1:100, Jackson Laboratory, Bar Harbor, USA) and rat serum (1:100, GeneTex, Irvine, USA) diluted in FACS buffer for 20 min at 4 °C to block of Fc receptors. Then, cells were incubated with LIVE/DEAD® Fixable Dead Cell Stain Kit (Thermo Fisher Scientific) diluted 1:1000, followed by incubation with fluorochrome-coupled antibodies (Supplementary Table S1) or corresponding isotype control antibodies diluted in FACS buffer for 30 min at 4 ° After fixation and permeabilization for 30 min at 4 °C using the Foxp3/Transcription Factor Staining Buffer Set (Thermo Fisher Scientific), resuspended cells were kept overnight at 4 °C. For intracellular stainings, fluorochrome-conjugated antibodies (diluted 1:100 in permeabilization buffer) were added to the cells, followed by incubation for 30 min at 4 °C. Samples were acquired at a BD FACS LSRFortessa Flow Cytometer (BD Biosciences) and analyzed with FlowJo V10 (Tree Star, Ashland, USA). Gating strategy is depicted in Fig. S10.

### Single-Cell RNA sequencing/Bulk sequencing

#### Library preparation

Single cells suspensions of eight digested tumors per group were pooled. Afterwards, cells were incubated with LIVE/DEAD® Fixable Dead Cell Stain Kit (Thermo Fisher Scientific) diluted 1:1000 in PBS for 30 min at 4 °C and stained with PE Rat Anti-Mouse CD45 antibody (BD) in FACS buffer for 30 min at 4 °C. Sorting for viable CD45^+^ cells was performed with a BD FACS Aria III.

The cell suspension of CD45 positive cells was adjusted to a concentration of approximately 1000 cells/μl. The cells were then loaded onto the 10x Genomics Chromium controller for droplet-enabled single cell RNA sequencing according to the manufacturer’s instruction manual. Library generation was performed following the Chromium Single Cell 3′ Reagents Kit version 3.1 user guide. The libraries were sequenced on the Illumina NovaSeq 6000 platform.

#### Pre-processing of individual scRNA experiments

Reads were aligned to the hg_GRCm38 reference genome with Cell Ranger v6.1.1 (10X genomics) using default parameters to produce cell-by-gene unique molecular identifier (UMI) count matrices. UMI count data was analyzed with Seurat R package v.4.0.5. Quality of datasets was assessed by UMI count, feature count and percentage of mitochondrial RNA reads. We filtered for high-quality cells based on feature count (above 1000) and percentage of mitochondrial reads (below 15%). Expression matrices of high-quality cells were log-normalized and scaled to a total expression of 1×10e5 UMIs for each cell. No regression of variables was applied.

#### Data Integration

2000 variable genes were determined per dataset using Seurat’s FindVariableFeatures function with “vst” selection method. To align the five libraries canonical correlation analysis (CCA) implementation was used in Seurat (Butler *et al*, 2018). Integration anchors were computed based on the five datasets using 50 dimensions and imputing 2000 integration genes of all variable genes. The resulting anchors were used to integrate the datasets.

#### Processing of integrated scRNA data

Integrated scRNA data was scaled and 50 principal components (PCs) were calculated. Next, Uniform Manifold Approximation and Projection (UMAP) was used for dimensionality reduction with 50 PCs, reducing to two dimensions. Nearest neighbors were calculated with 50 PCs and several clustering resolutions were computed with Seurat’s FindClusters function. After manual inspection, resolution 0.2 was used for analysis of the dataset including all lineages, resulting in 13 clusters. Differential expression analysis was performed using Seurat’s FindAllMarkers function on the clustering resolution 0.2.

The contribution of the different treatment groups to the clusters was assessed by calculating the percentage of a respective cluster within the treatment group, to account for differences in library size. The clustering was plotted using DimPlot and individual gene UMAP plots were generated using the FeaturePlot function with minimum expression of q01 and maximum expression of q99.

#### Processing of myeloid cells sub-clustering

The main cluster of cells was myeloid cells. To improve resolution, this group was analyzed separately by extracting clusters 0, 1, 3, 4, and 9 of resolution 0.2. pDCs were not included in the sub-clustering as they clustered far away from other myeloid cells. On the myeloid subset the UMAP coordinates, nearest neighbors and clusters were re-calculated using 50 PCs. After manual inspection, resolution 0.5 was used for further analysis resulting in 13 different clusters. The contribution of the different treatment groups to the clusters was assessed by calculating the percentage of a respective cluster within the treatment group to account for differences in library size. Differential expression analysis was performed on the resolution 0.5. Plotting of clustering and genes was performed as outlined above.

#### Processing of lymphoid cells sub-clustering

The second largest group of cells was lymphoid cells. To improve resolution, this group was analyzed separately by extracting clusters 2, 5, 6, 8, and 10 of resolution 0.2. On the lymphoid subset the UMAP coordinates, nearest neighbors and clusters were re-calculated using 50 PCs. After manual inspection resolution 0.25 was used for further analysis, resulting in 10 different clusters. The contribution of the different treatment groups to the clusters was assessed by calculating the percentage of a respective cluster within the treatment group to account for differences in library size. Differential expression analysis was performed on the resolution 0.25. Plotting of clustering and genes was performed as outlined above.

#### Differential Expression heatmaps per cluster

To ensure proper cell type annotation we used the results of the differential expression analysis, as well as correlation to reference datasets (see below). Based on the differential expression analysis the enriched genes were selected: for the heatmap of all cells (Fig. 3B) all genes with average log2 fold-change (avg-log2FC) > 0.6 and adjusted p-value (p-adj.) < 1e-100 were selected. Additionally, mitochondrial, and ribosomal genes were excluded to reduce the number of genes in the heatmap. For myeloid (Fig. 4C) and lymphoid cells (Fig. 5C) all genes with avg-log2FC > 0.6 and p-adj. < 1e-50 were selected. For each respective dataset the average expression per cluster was calculated for the selected genes using Seurat’s AverageExpression function. The order of clusters (columns) in the heatmap was manually selected. Per cluster, the genes were ordered by ascending p-adj. and descending avg-log2FC. Genes enriched in multiple clusters were only kept in the cluster with lowest p-adj. and highest avg-log2FC resulting in unique rows. To plot the table as a heatmap the pheatmap v1.0.12 package was used in R. Expression was scaled per gene and individual genes were selected for annotation. The color of the gene annotation reflects the cluster each gene was found to be enriched in based on the criteria outlined above.

The entirely ordered table of genes for each heatmap as well as the differential gene expression results are reported in Supplementary Tables S2 (all cells), S3 (myeloid cells), and S7 (lymphoid cells).

#### Correlation to references for myeloid cells

To support cell type annotation, we compared our scRNA data to published datasets. For comparison of myeloid cells, we downloaded two datasets: Gubin et al. (Gubin *et al*., 2018) and Zhang et al. (Zhang *et al*., 2020). Processed read matrices and metadata were downloaded from GEO (accession number: GSE119352) and ArrayExpress (accession number: PRJEB34105), respectively. The data of Gubin et al. was filtered as outlined in the Methods part by the authors. To perform correlation to our data we only selected the anti-CTLA4 and control group. For correlation, datasets were transferred to monocle3 v1.0.0 (Trapnell *et al*., 2014). Count and cell metadata were used to construct a cell data set object. For clustering, the tSNE coordinates provided in the metadata were used. Next, only myeloid cells were selected, and gene modules were calculated using the monocle’s find_gene_modules function on the reference dataset. This resulted in 32 modules including approximately 2000 genes. Myeloid cell sub-clustering of our data was transferred to monocle as outlined above, followed by aggregation of gene expression per module and cell type. For our Myeloid Cells we used the resolution as presented above and for Macrophages of Gubin et al. (Gubin *et al*., 2018) we used the cell type annotation provided in the metadata. Pearson correlation was performed between the different clusters and plotted using the corrplot v0.92 package in R. Gene modules, aggregated expression per cluster and the correlation matrix are reported in Supplementary Table S4.

The same approach was used to compare to the myeloid cells in Zhang et al. (Zhang *et al*., 2020). Only the data of the aCSF1R experiment (isotype and treatment) were used. The data was filtered as reported in the Methods part, and UMAP coordinates of the sub-clustering provided by the metadata were used. The data was transferred to monocle3 as outlined above, and gene modules were calculated resulting in 37 modules including approximately 2000 genes. Gene expression was aggregated per module and sub-cluster for both reference and our data and Pearson correlation was plotted as described above. Correlation plots were generated for Macrophages/Monocytes and DCs. To report the full spectrum of DCs in our dataset, the correlation of DCs included the cDC clusters of the sub-clustering and the pDC cluster that is only found in Fig. 3. Gene modules, aggregated expression per cluster and the correlation matrix are reported in Supplementary Table S5.

#### Correlation to reference for lymphoid cells

To support the cell type annotation of the lymphoid cell sub-clustering we used a recently published dataset of TILs (Andreatta *et al*., 2021). The TIL atlas data was downloaded from figshare as provided by the authors GitHub page (https://github.com/carmonalab/ProjecTILs). The same approach as outlined for myeloid cells was used to correlate the lymphoid clusters. As our data included cell types that were not present in the lymphoid atlas (e.g. NK cells), we calculated gene modules on our data and then correlated to the reference. This resulted in 34 modules of 2000 genes. Plotting was performed as outlined above. Gene modules, aggregated expression per cluster and the correlation matrix are reported in Supplementary Table S8.

#### Differential Expression Analysis and GO-terms for myeloid cells

To compare the different treatment groups within macrophages and DCs we performed pairwise differential expression analysis using Seurat’s FindMarkers function. Comparing irradiated (C12-iF and C12/aC4-iF) with untreated tumors revealed a lot of DEGs, both per cluster and comparing all myeloid cells were detected. The DEGs comparing all myeloid cells were plotted using the EnhancedVolcano v1.10 package in R with log2FC > ±0.5. The result of differential expression analysis for myeloid cells is reported in Supplementary Table S6. Similarly, C12/anti-CTLA4-treated (iF and oF combined) were compared with C12 treated (iF and oF), aggregating all TAM clusters (TAM-1 - TAM-7) and plotted as outlined previously.

Enriched GO-terms among DEGs were determined performing gene set enrichment analysis (GSEA) with the clusterProfiler v4.0.5 package in R. DEGs were defined with p-adj. <0.05 and avg-log2FC > 0.5. Pairwise similarity of enriched GO-terms was assessed using the pairwise_termsim function, and hierarchical clustering of GO-terms was plotted. The clusters of GO-terms were annotated manually.

To test the enrichment of different GO-terms per group and cell type we generated gene set scores for a selection of GO-terms. The full list of genes per GO-term was downloaded using the biomaRt v2.48.3 package in R. Seurat’s AddModuleScore function was used to assign scores per GO-term to each cell. For each treatment group and cell type the median score was calculated, scaled, and plotted using the pheatmap package in R. This was performed for: 1. only macrophage and monocyte clusters, 2. all clusters of myeloid sub-clustering, 3. all clusters of Fig. 3 that are not included in myeloid sub-clustering (Fig. 4), 4. aggregated for all macrophages and monocyte clusters per treatment group and the aggregated macrophages of the Gubin et al. paper.

#### Differential Expression Analysis and GO-terms for lymphoid cells

Within lymphoid cells we focused on comparisons within the out-of-field tumors, as they made up most of the T cell clusters. We performed pairwise differential expression analysis comparing C12-oF and C12/aC4-oF lymphoid cells per cluster using Seurat’s FindMarkers function. No strongly enriched genes (defined as p-adj. <0.05 and avg-log2FC > 1) were found in activated clusters, thus we focused on the naïve cells. Enriched GO-terms (on DEGs defined as p-adj. <0.05 and avg-log2FC > 1) and GO-scores were assessed as described above. In the scores we included GO-terms of the GSEA analysis and additional GO-terms describing important T cell functions. The result of differential expression analysis for lymphoid cells is reported in Supplementary Table S9.

#### Comparison of FACS and scRNA data

To compare cell type composition in pooled scRNA libraries with individual flow cytometry experiments we used a recently published scoring method (Andreatta *et al*., 2021) (ref https://www.sciencedirect.com/science/article/pii/S2001037021002816?via%3Dihub). The genes of proteins used for flow cytometric phenotyping were aggregated to a rank-based score. The percentage of cells positive for all the markers included in the score (passing the score with >0.25 for one marker, >0.5 for two markers, and >0.66 for three markers) was used to compare to the percentages of positive cells determined by flow cytometry. Comparability was ensured as both flow cytometry and scRNA data were obtained from the same CD45^+^ starting population.

Following markers were used to generate scores:

NK: Klrb1c; CD8: Cd8a + Cd8b1; CD8 LAG3: Cd8a + Cd8b1 + Lag3, CD8 PD1: Cd8a + Cd8b1 + Pdcd1; CD8 CD25: Cd8a + Cd8b1 + Il2ra; CD8 CTLA4: Cd8a + Cd8b1 + Ctla4; CD8 CD69: Cd8a + Cd8b1 + Cd69; CD8 NT5E: Cd8a + Cd8b1 + Nt5e; CD4: Cd4; CD4 NT5E: Cd4 + Nt5e; CD4 CTLA4: Cd4 + Ctla4; CD4 CD69: Cd4 + Cd69; CD4 PD1: Cd4 + Pdcd1; CD4 LAG3: Cd4 + Lag3 For Tregs and DCs we used the cluster percentage to compare to flow cytometry.

As TAM clusters formed a dynamic continuum in the scRNA data, we did not generate scores for comparison with flow cytometry experiments, but compared groups within the two modalities separately.

#### RNA velocity analysis

Quantification of spliced and unspliced counts per cell was performed using velocyto v0.17.17 using CellRanger aligned bam files as input, with default settings and masking repeat elements. The generated loom files were further processed in python with jupyter notebook v6.4.3 using scvelo v0.2.3, scanpy v1.8.1, anndata v0.7.6, numpy v1.21.1-concatenating datasets, performing scvelo analysis with default settings including preprocessing, computation of moments of spliced/unspliced abundances, velocity estimation using a stochastic model of transcriptional dynamics with differential kinetic estimation and velocity graph computation, followed by projecting velocities into the low dimensional UMAP embedding. Seurat objects with UMAP coordinates and annotations were transferred to scvelo by exporting them as loom files using as.loom of SeuratDisk v0.0.09019. Trajectories were plotted in UMAP projection either containing all cells or showing individual treatment groups.

### Tumor re-challenge and analysis of tumor-specific immune responses

Mice showing complete tumor rejection upon treatment were re-challenged 3 to 6 months later by s.c. injection of 4 × 10^5^ EO771cells into the right flank. Naïve (untreated) mice were used as controls and tumor growth of all mice was monitored. After 14 days, spleens were aseptically removed and passed through a 70 µm cell strainer to obtain single cell suspensions. Erythrocytes were lysed using eBioscience(tm) 1X RBC Lysis Buffer (Thermo Fischer Scientific). Splenocytes were resuspended in T cell medium consisting of alpha MEM (Sigma-Aldrich) supplemented with 10% heat-inactivated FCS, 2 mM L-glutamine (Thermo Fischer Scientific), 50 µM 2-mercaptoethanol (Thermo Fischer Scientific), 2.5% (v/v) supernatant of concanavalin A stimulated rat spleen cell cultures, 12.5 mM methyl-α-d-mannopyranoside (Thermo Fisher Scientific), 100 U/ml penicillin, 100 μg/ml streptomycin. EO771, B16F10, and RMA cells were harvested in T cell medium and irradiated with 250 Gy. Briefly, 1 × 10^6^ splenocytes were co-cultured with 5 × 10^4^ target cells in triplicates and incubated for 20 h.

### Statistical analysis

Statistical analysis was calculated using GraphPad Prism 7.05 (GraphPad Software, San Diego, USA). The respective statistical test is stated in the figure legends. Levels of significance are defined as follows:* p ≤ 0.05, ** p ≤ 0.01; *** p ≤ 0.001: **** p ≤ 0.0001). Further statistical analysis was conducted using R 4.0.5 (https://www.R-project.org/) and the R package lmerTest (Kuznetsova *et al*, 2017). In this analysis the effects of the treatment groups (C4, C12-aC4, C12 and untreated) and the tumor location (in-field and out-field) on various outcome parameters was investigated using linear mixed models. The primary motivation for using this class of models stems from the dependence between tumor samples taken from the same mouse. Linear mixed models are an extension of simple linear models, which incorporates both fixed effects (group and tumor location) and random intercept (mice). Results of this analysis are given in detail in Supplementary Table S10, all percentages of the FACS results are shown in Supplementary Table S11.

## Acknowledgements

We thank Lena Vogelbacher, Nora Schuhmacher and Claudia Rittmüller for their excellent technical assistance. We are grateful to the DKFZ single cell lab and High Throughput Sequencing Unit for scRNA sequencing. Furthermore, we thank the Division of Biostatistics of the DKFZ, in particular Tim Holland-Letz, for their support in biostatistic analysis. We also acknowledge the help of the Flow Cytometry Core Facility and thank the Center for Preclinical Research at the DKFZ for excellent animal care take. Moreover, we thank Michael Baumann and Ina Kurth from the Division of Radiooncology / Radiobiology for providing the Faxitron MultiRad225 irradiator as well as Armin Runz and Peter Haering (DKFZ) for their help in handling the irradiator device.

This study was supported in part by Zentrum für Personalisierte Medizin (ZPM-Network BW, Project PROMI), collaborative research center of the German Research Foundation (DFG, Unite, SFB-1389, Project 404521405), and intramural funds of the National Center for Tumor Disease (NCT) and the German Cancer Consortium (DKTK) radiation oncology programs.

## Author contributions

Conceptualization: SBE, SR, WO, LH, JD, AA, MM

Methodology: LH, WO, MM, AA, SB

Investigation: LH, ED, MM, JF

Visualization: LH, OLE, TK, CR

Supervision: SBE, SR, WO

Writing—original draft: LH, WO, OLE, SBE

Writing—review & editing: LH, OLE, WO, SBE, TK, AA

## The Paper Explained

### Problem

Radiotherapy can control local tumor growth and may have immunostimulatory capacity. In order to prevent outgrowth of metastasis, current immunotherapy approaches are based on irradiation combined with immune checkpoint inhibition (radioimmunotherapy). Carbon ion radiotherapy shows superior biophysical properties over conventional photon irradiation, however, studies on the immunological effects induced by carbon ion irradiation (C12) versus photon irradiation combined with different checkpoint inhibitors (anti-CTLA4 or anti-PD-L1) are lacking. Furthermore, differential gene expression profiling on specific immune cell subpopulations accumulating in irradiated versus non-irradiated tumors induced by such treatment regimens are missing.

### Results

We have established a preclinical mouse model allowing in depth analysis of different combination therapies using irradiation and immune checkpoint inhibitors. Our study demonstrates superior effects of carbon ion-based radiotherapy combined with CTLA4 blockade over PD-L1 inhibition. This regimen eradicated irradiated tumors and of note, mediated abscopal rejection of non-irradiated tumors. Remarkably, direct irradiation induced conversion of the intratumoral myeloid compartment and accumulation of NK cells, while non-irradiated tumors showed an increase in activated CD8+ T cells upon co-application of CTLA4 blocking antibody.

### Impact

Showing that carbon ion radiotherapy is eligible for combination with CTLA4 blockade, our study is of remarkable translational interest. According to the results presented, validation of novel clinical radioimmunotherapy approaches should not only include a thorough investigation of the immunological responses induced at the irradiation site, but also of local immune reactions in non-irradiated, distant metastases.

### Data Availability Section

All raw and processed sequencing data generated in this study will be deposited and shared upon publication and accession numbers provided. All custom code used in this study will be made available upon acceptance of the paper under GitHub Link and DOI.

