## Supplemental figures S1-S10 and Suppl Table S1 for "Carbon ion irradiation plus CTLA4 blockade elicits therapeutic immune responses in a murine tumor model"

**This PDF file includes:**

Figs. S1 to S10
Table S1
References of the Supplemental Material

**Other Supplementary Materials for this manuscript include the following:**
These excel files are provided as separate supplemental files.

Table S2: Data for heatmap and DEGs all cells (Fig. 3)
Table S3: Data for heatmap and DEGs myeloid cells (Fig. 4)
Table S4: Modules for comparison of myeloid cells to Zhang et al.
Table S5: Modules for comparison of myeloid cells to Gubin et al.
Table S6: DEGs pairwise in myeloid cells of C12-iF and C12/aC4-iF vs untreated
Table S7: Data for heatmap and DEGs lymphoid cells (Fig. 5)
Table S8: Modules for comparison of lymphoid cells to Andreatta et al.
Table S9: DEGs pairwise in lymphoid cells of C12-oF vs C12/aC4-oF
Table S10: Statistics mixed model for FACS analysis
Table S11: FACS percentages

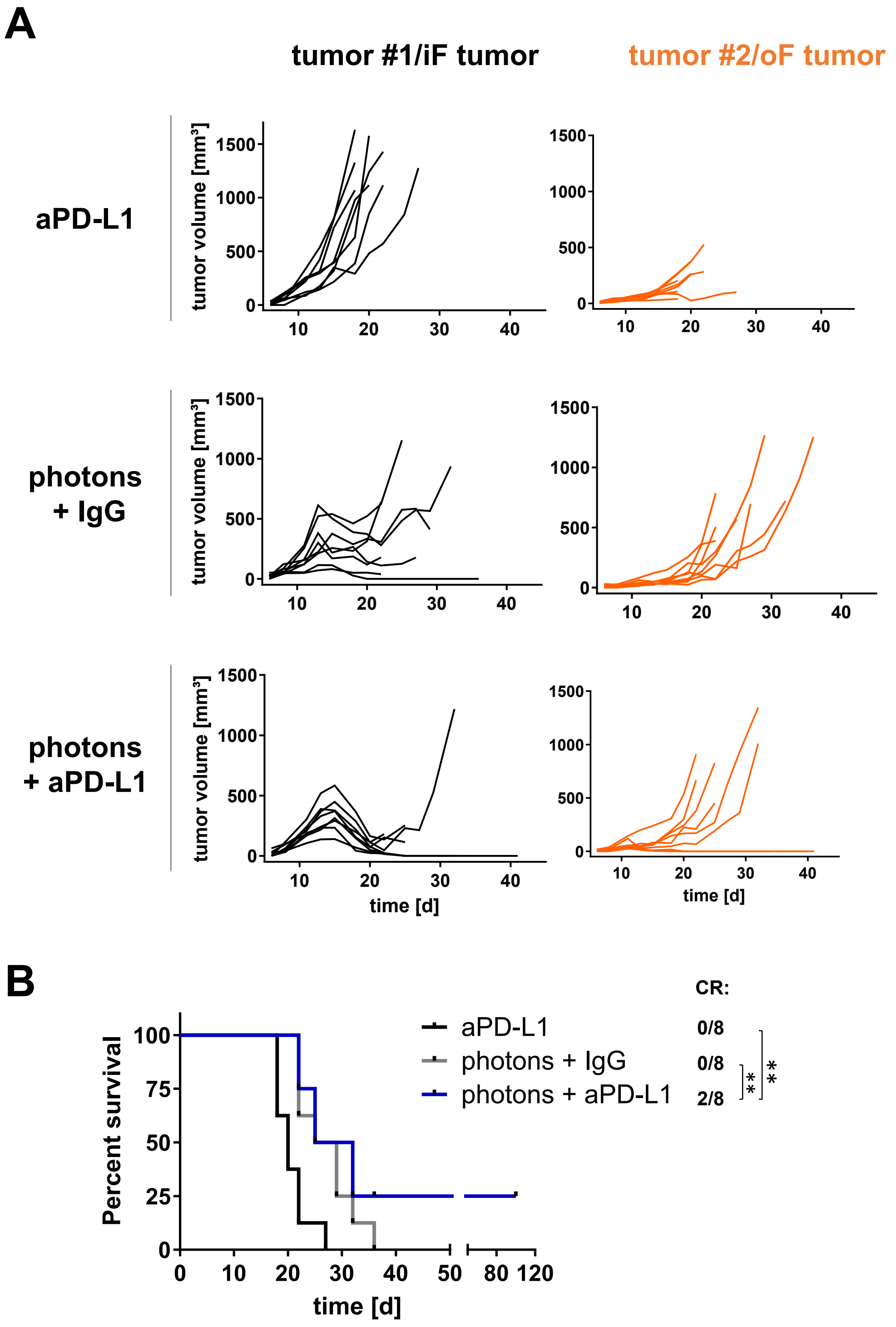


**Fig. S1. Combination of photon RT with checkpoint blockade against PD-L1.**
(A) Individual growth curves of tumor #1/in -field (iF) tumors and tumor #2/out-of-field (oF) tumors of mice treated with anti-PD-L1 antibody only (upper panel), photon irradiation (middle panel), or combination of photon irradiation and anti-PD-L1 antibody.
(B) Kaplan-Meier survival analysis. In addition, complete response (CR) rates for each group are shown. Significance was determined using a log-rank (Mantel-Cox) test with Holm-Bonferroni correction.

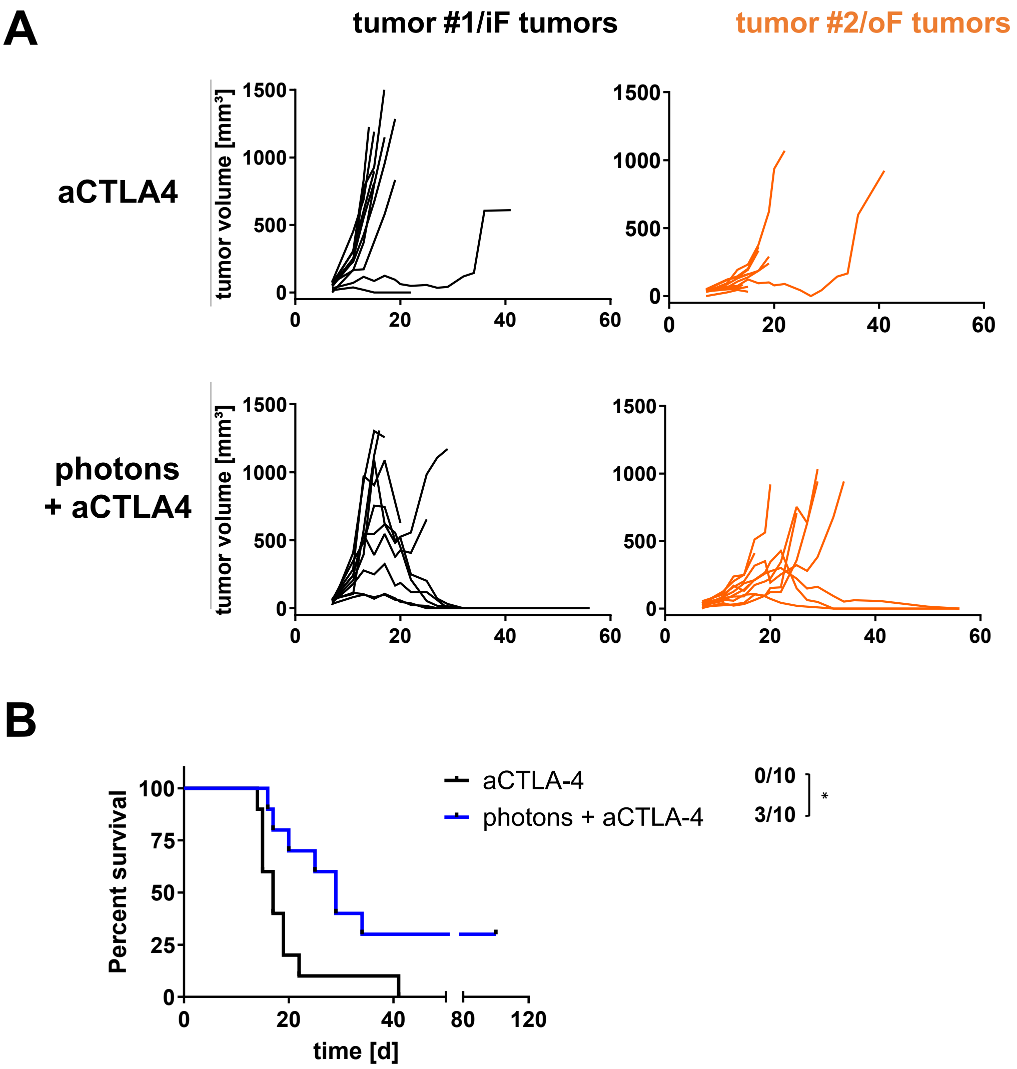


**Fig. S2. Combining photon RT with checkpoint blockade against anti-CTLA4.**

**(A)** Individual growth curves of tumor #1 and tumor #2 of mice treated as indicated.

**(B)** Kaplan-Meier survival analysis. In addition, complete response rates for each group are shown. Significance was determined using a log-rank (Mantel-Cox) test with Holm-Bonferroni correction.

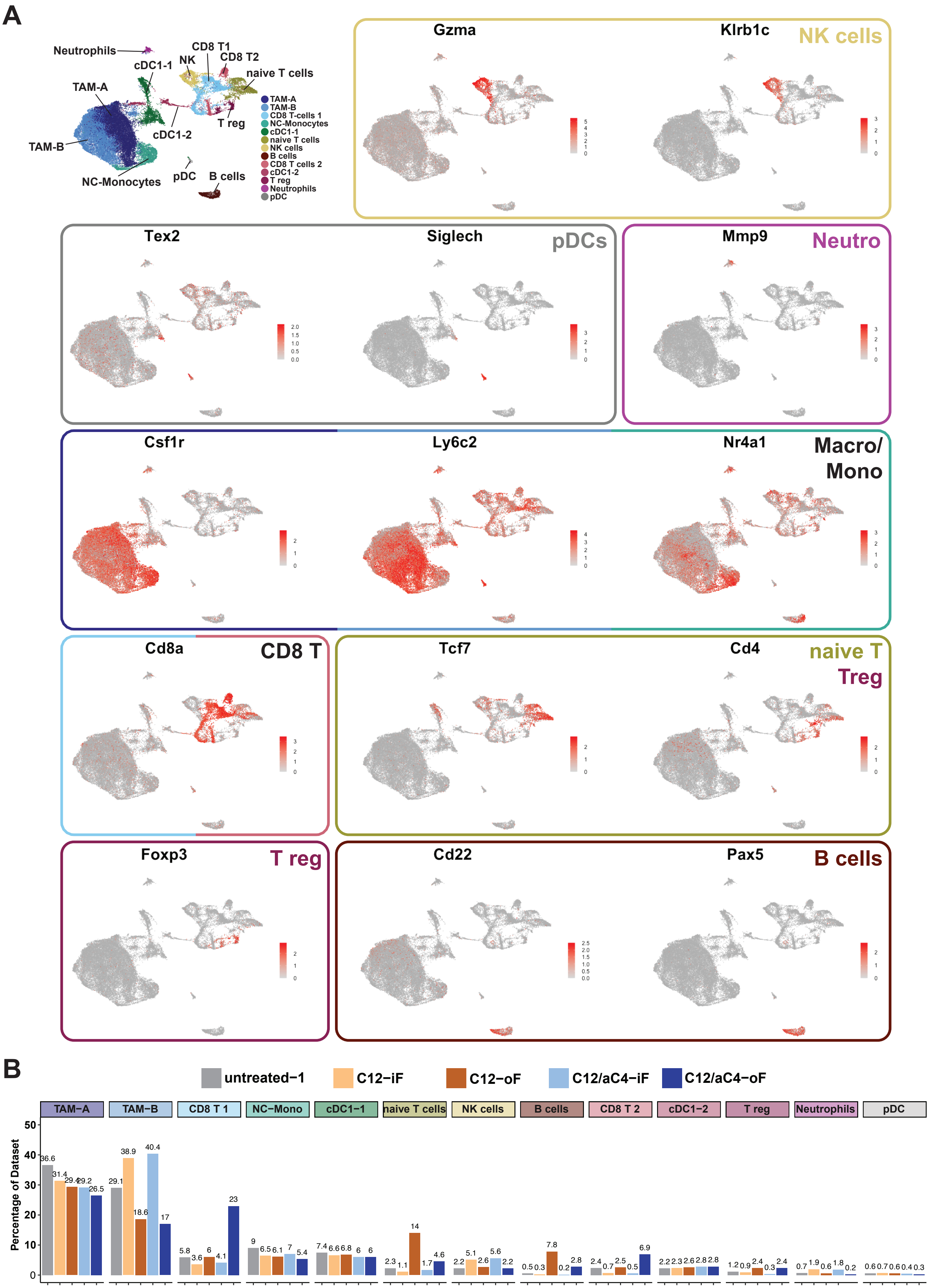
A
Neutrophils
cDC1-1
NK
CD8 T1
CD8 T2
naive T cells
TAM-A
TAM-B
cDC1-2
T reg
NC-Monocytes
pDC
B cells
TAM-A
TAM-B
CD8 T-cells 1
NC-Monocytes
cDC1-1
naive T cells
NK cells
B cells
CD8 T cells 2
cDC1-2
T reg
Neutrophils
pDC
Gzma
Klrb1c
NK cells
Tex2
Siglech
pDCs
Mmp9
Neutro
Csf1r
Ly6c2
Nr4a1
Macro/ Mono
Cd8a
CD8 T
Tcf7
Cd4
naive T
Treg
Foxp3
T reg
Cd22
Pax5
B cells
B
untreated−1
C12−iF
C12−oF
C12/aC4−iF
C12/aC4−oF
TAM−A
TAM−B
CD8 T 1
NC−Mono
cDC1−1
naive T cells
NK cells
B cells
CD8 T 2
cDC1−2
T reg
Neutrophils
pDC
Percentage of Dataset
0
10
20
30
40
50
36.6 31.4 29.4 29.2 26.5
29.1 38.9 18.6 40.4 17
5.8 3.6 6 4.1 23
9 6.5 6.1 7 5.4
7.4 6.6 6.8 6 6
2.3 1.1 14 1.7 4.6
2.2 5.1 2.6 5.6 2.2
0.5 0.3 7.8 0.2 2.8
2.4 0.7 2.5 0.5 6.9
2.2 2.3 2.6 2.8 2.8
1.2 0.9 2.4 0.3 2.4
0.7 1.9 0.6 1.8 0.2
0.6 0.7 0.6 0.4 0.3

**Fig. S3. Overview clustering reveals immune infiltrating subtypes.**

(**A**) Markers of cell types shown in Fig. 3B and D. Upper left: clustering reference of the of tumor infiltrating immune cell spectrum. Established markers identify NK cells (Gzma, Klrb1c = Nk1.1), pDCs (Tex2, Siglech), Neutrophils (Mmp9), Macrophages/Monocytes (Csf1r, Ly6c2, Nr4a1), CD8 T cells (Cd8a), naïve T cells (Tcf7, Cd4), regulatory T cells (Foxp3), and B cells (Cd22, Pax5).

(**B**) Cluster contribution of different treatment groups. Cluster annotation corresponds to Fig. 3B.

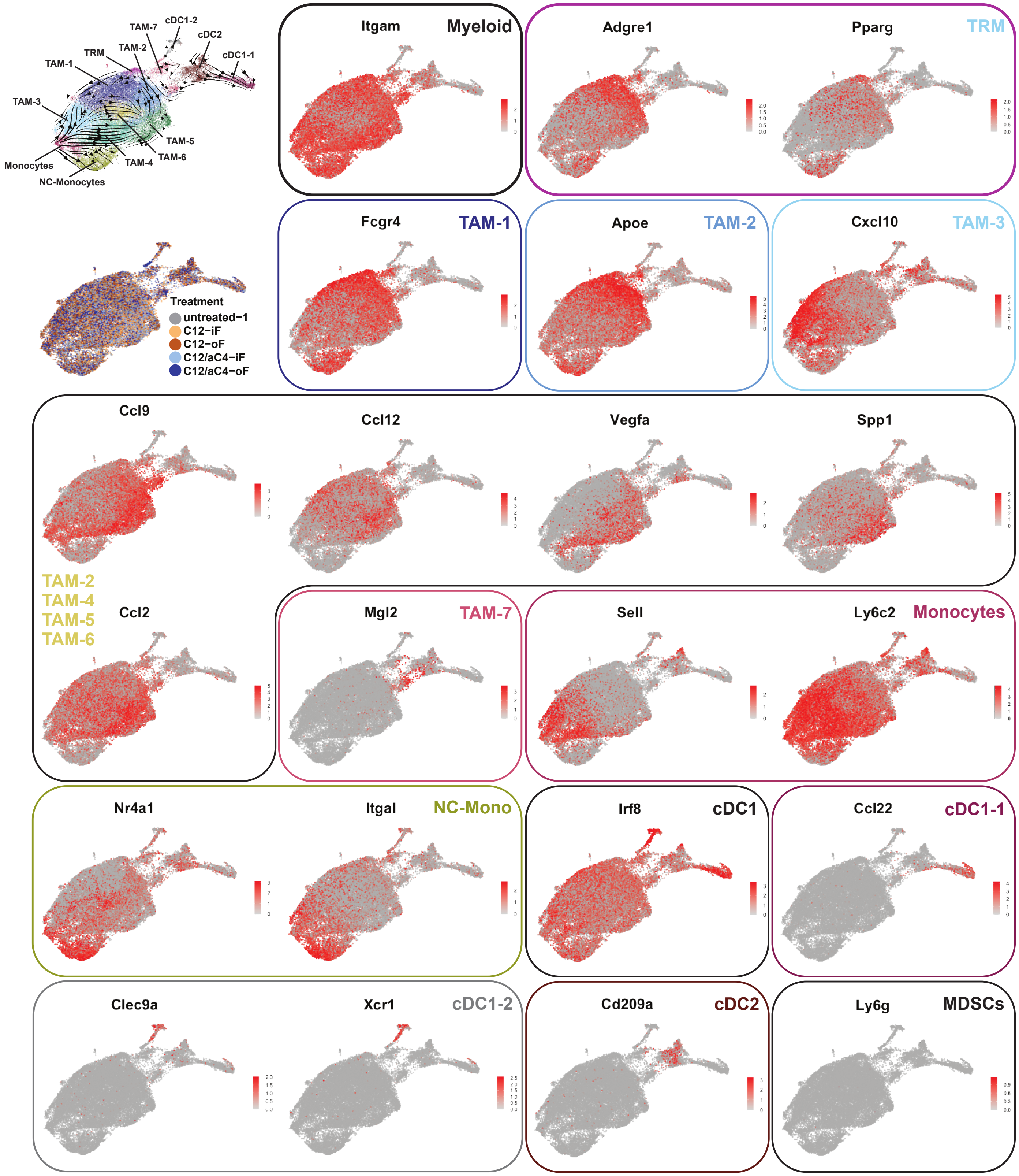


**Fig. S4. Marker genes for myeloid subclustering.** UMAPs of selected genes for myeloid cell subclustering. The two left top graphs show clustering and velocity as in Fig. 4A (upper graph), as well as clustering colored by treatment groups (lower graph).

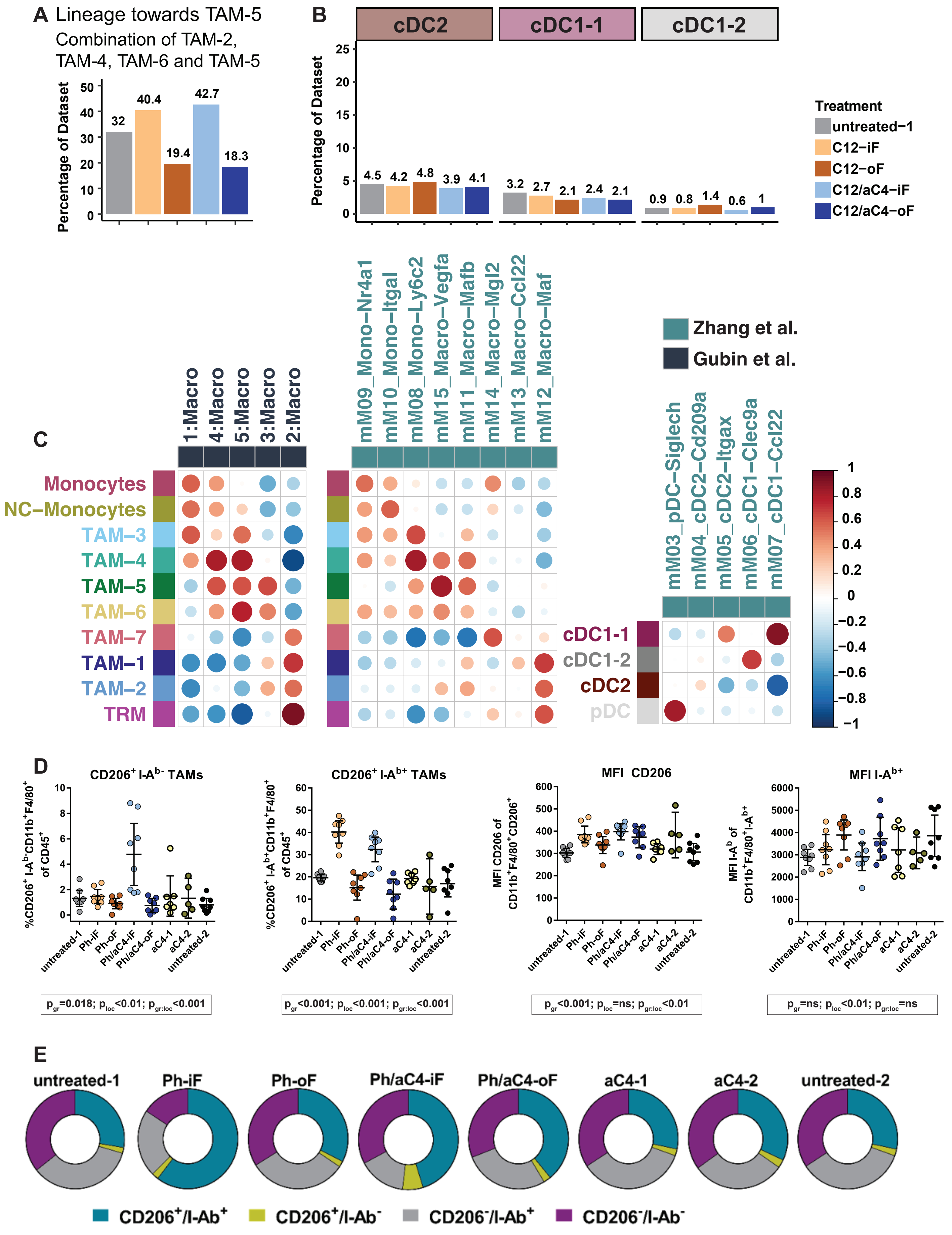
A Lineage towards TAM-5
Combination of TAM-2, TAM-4, TAM-6 and TAM-5
Percentage of Dataset
32
40.4
19.4
42.7
18.3
B
cDC2
cDC1-1
cDC1-2
Percentage of Dataset
4.5 4.2 4.8 3.9 4.1
3.2 2.7 2.1 2.4 2.1
0.9 0.8 1.4 0.6 1
Treatment
untreated−1
C12−iF
C12−oF
C12/aC4−iF
C12/aC4−oF
C
1:Macro
4:Macro
5:Macro
3:Macro
2:Macro
mM09_Mono−Nr4a1
mM10_Mono−Itgal
mM08_Mono−Ly6c2
mM15_Macro−Vegfa
mM11_Macro−Mafb
mM14_Macro−Mgl2
mM13_Macro−Ccl22
mM12_Macro−Maf
Zhang et al.
Gubin et al.
mM03_pDC−Siglech
mM04_cDC2−Cd209a
mM05_cDC2−Itgax
mM06_cDC1−Clec9a
mM07_cDC1−Ccl22
Monocytes
NC−Monocytes
TAM−3
TAM−4
TAM−5
TAM−6
TAM−7
TAM−1
TAM−2
TRM
cDC1-1
cDC1-2
cDC2
pDC
D
CD206+ I-Ab- TAMs
%CD206+ I-Ab-CD11b+F4/80+ of CD45+
CD206+ I-Ab+ TAMs
%CD206+ I-Ab+CD11b+F4/80+ of CD45+
MFI CD206
MFI CD206 of CD11b+F4/80+CD206+
MFI I-Ab+
MFI I-Ab of CD11b+F4/80+I-Ab+
untreated-1
Ph-iF
Ph-oF
Ph/aC4-iF
Ph/aC4-oF
aC4-1
aC4-2
untreated-2
pgr=0.018; ploc<0.01; pgr:loc<0.001
pgr<0.001; ploc<0.001; pgr:loc<0.001
pgr<0.001; ploc=ns; pgr:loc<0.01
pgr=ns; ploc<0.01; pgr:loc=ns
E
untreated-1
Ph-iF
Ph-oF
Ph/aC4-iF
Ph/aC4-oF
aC4-1
aC4-2
untreated-2
CD206+/I-Ab+
CD206+/I-Ab-
CD206-/I-Ab+
CD206-/I-Ab-

**Fig. S5. Comparison to published data reveals identity of increased TAM populations.**

**(A)** Aggregated percentages of all clusters along the lineage towards TAM-5 (TAM-2, -4, -6 and TAM-5) reveals increase in iF tumors.

**(B)** Percentages of cDC clusters in different treatment groups.

(**C**) Correlation heatmaps of subclustered myeloid clusters of this paper with two reference datasets (Gubin *et al*, 2018; Zhang *et al*, 2020). Clusters were arranged following the trajectories from Monocytes (from top) and from TRM (from bottom) towards differentiation end points. Modules of co-expressed genes were calculated on the reference datasets (see Materials and Methods) and gene expression was aggregated per cluster and module. Aggregated expression values were used for correlation comparing Macrophages/Monocytes of this paper to either Gubin et al. (Gubin *et al.*, 2018) or Zhang et al. (Zhang *et al.*, 2020) or comparing DCs of this paper to Zhang et al. (Zhang *et al.*, 2020). pDCs were added from the clustering in Fig. 3 to include all DC groups. Color code and dot size show pairwise correlation from anti-correlation (-1, dark blue and large point) over no correlation (0, white and small point) to high correlation (1, red and large point). Gene modules as well as aggregated expression values can be found in **Table S4** and **Table S5**.

(**D**) UMAP depicting expression of Folr2 in myeloid cells is enriched in TAM-2 and TAM-5 groups.

(**E**) and (**F**) Proportion of infiltrating immune cell subpopulations within individual tumors determined by flow cytometry (E) or summarized as pie charts (F) following photon-based treatment schedules depicted in Fig. 1A.

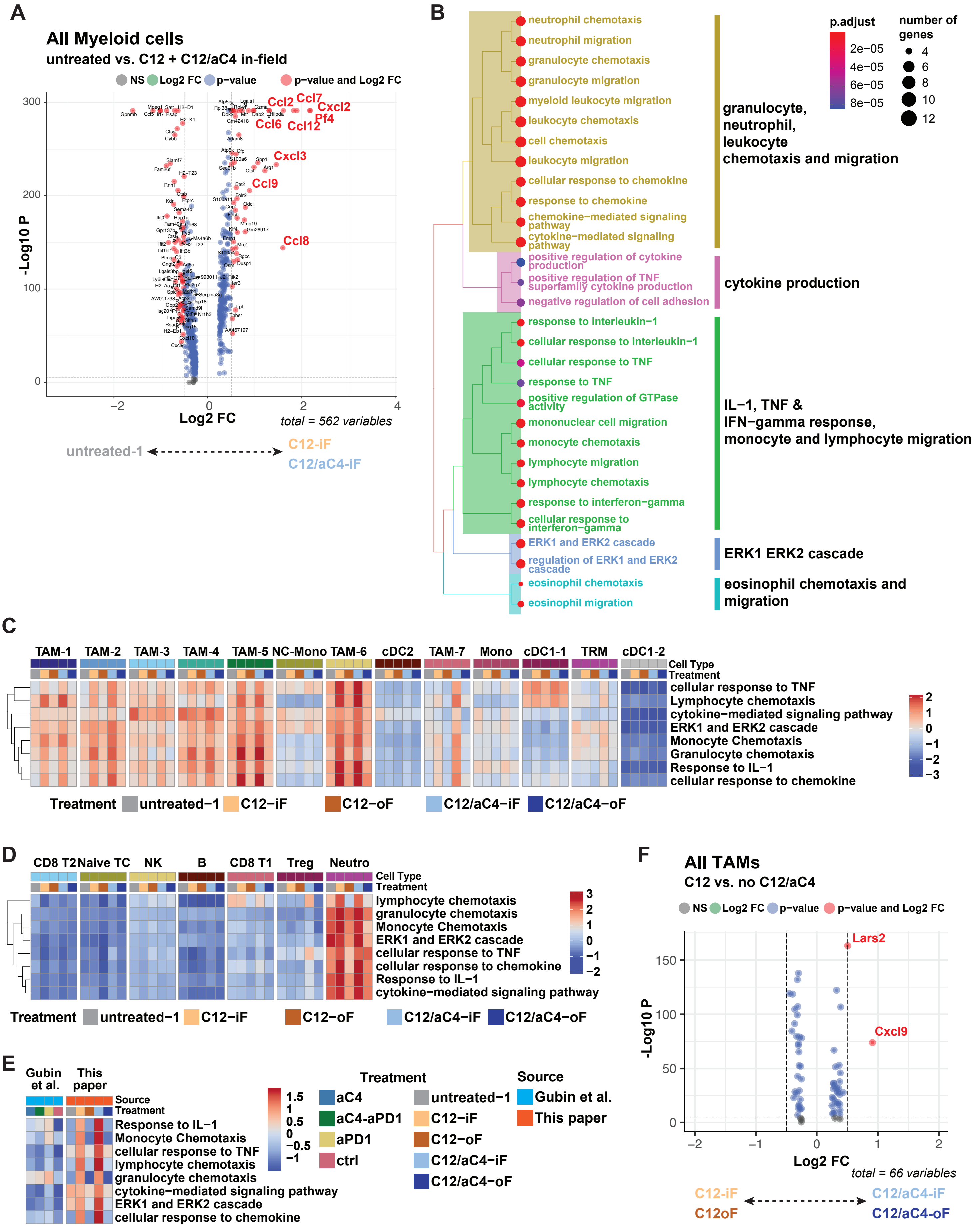

A
All Myeloid cells
untreated vs. C12 + C12/aC4 in-field
NS Log2 FC p−value p−value and Log2 FC
-Log10 P
Log2 FC
total = 562 variables
Ccl2 Ccl7 Cxcl2 Pf4 Ccl6 Ccl12 Cxcl3 Ccl9 Ccl8
untreated-1
C12-iF
C12/aC4-iF
B
neutrophil chemotaxis
neutrophil migration
granulocyte chemotaxis
granulocyte migration
myeloid leukocyte migration
leukocyte chemotaxis
cell chemotaxis
leukocyte migration
cellular response to chemokine
response to chemokine
chemokine−mediated signaling pathway
cytokine−mediated signaling pathway
positive regulation of cytokine production
positive regulation of TNF superfamily cytokine production
negative regulation of cell adhesion
response to interleukin−1
cellular response to interleukin−1
cellular response to TNF
response to TNF
positive regulation of GTPase activity
mononuclear cell migration
monocyte chemotaxis
lymphocyte migration
lymphocyte chemotaxis
response to interferon−gamma
cellular response to interferon−gamma
ERK1 and ERK2 cascade
regulation of ERK1 and ERK2 cascade
eosinophil chemotaxis
eosinophil migration
granulocyte, neutrophil, leukocyte chemotaxis and migration
cytokine production
IL−1, TNF & IFN−gamma response, monocyte and lymphocyte migration
ERK1 ERK2 cascade
eosinophil chemotaxis and migration
p.adjust
2e−05
4e−05
6e−05
8e−05
number of genes
4
6
8
10
12
C
TAM-1 TAM-2 TAM-3 TAM-4 TAM-5 NC-Mono TAM-6 cDC2 TAM-7 Mono cDC1-1 TRM cDC1-2
Cell Type
Treatment
cellular response to TNF
Lymphocyte chemotaxis
cytokine−mediated signaling pathway
ERK1 and ERK2 cascade
Monocyte Chemotaxis
Granulocyte chemotaxis
Response to IL−1
cellular response to chemokine
Treatment untreated−1 C12−iF C12−oF C12/aC4−iF C12/aC4−oF
D
CD8 T2 Naive TC NK B CD8 T1 Treg Neutro
Cell Type
Treatment
lymphocyte chemotaxis
granulocyte chemotaxis
Monocyte Chemotaxis
ERK1 and ERK2 cascade
cellular response to TNF
cellular response to chemokine
Response to IL−1
cytokine−mediated signaling pathway
Treatment untreated−1 C12−iF C12−oF C12/aC4−iF C12/aC4−oF
E
Gubin et al.
This paper
Source
Treatment
Response to IL−1
Monocyte Chemotaxis
cellular response to TNF
lymphocyte chemotaxis
granulocyte chemotaxis
cytokine−mediated signaling pathway
ERK1 and ERK2 cascade
cellular response to chemokine
Treatment
aC4
aC4-aPD1
aPD1
ctrl
untreated−1
C12−iF
C12−oF
C12/aC4−iF
C12/aC4−oF
Source
Gubin et al.
This paper
F
All TAMs
C12 vs. no C12/aC4
NS Log2 FC p−value p−value and Log2 FC
-Log10 P
Log2 FC
total = 66 variables
Lars2
Cxcl9
C12-iF
C12oF
C12/aC4-iF
C12/aC4-oF

**Fig. S6. Direct C12 radiotherapy induces strong gene expression changes in TAM subsets.**

**(A)** Differential gene expression analysis comparing myeloid cells from in field tumors following radiomonotherapy (C12-iF) or radioimmunotherapy (C12/aC4 iF). All Macrophages/Monocytes and DCs were aggregated to investigate overall effects of direct radiation. All genes passing a p-value of 0.05 and a log2 fold-change of 0.5 are annotated. Selected chemokines induced by direct radiation are depicted in red. List of differentially expressed genes including differential expression of individual clusters is reported in **Table S6**.

**(B)** Gene set enrichment analysis (GSEA) of genes enriched upon direct irradiation (A). GO-terms are clustered according to similarity to derive functional groups (see Materials and Methods). Adjusted p-value is shown from blue to red and the number of genes contributing to the respective score are depicted by dot size.

(**C**) Aggregated expression of selected enriched GO-terms of (B) per treatment group and cell type across all clusters of myeloid subclustering (related to Fig. 4F).

(**D**) Aggregated expression of selected enriched GO-terms of (B) per treatment group and cell type across all clusters of overview clustering in Fig. 3B that are not included in the myeloid subclustering.

(**E**) Comparison of macrophages (all clusters of myeloid subclustering except for cDCs) of this work with different immunotherapy treatment groups presented by Gubin et al. (Gubin *et al.*, 2018) reveals induction of enriched GO-terms is specific to in-field radiation.

(**F**) Differential gene expression analysis testing the additive effect of anti-CTLA4 treatment to C12 radiotherapy. Radiomonotherapy applications (C12-iF and -oF) were compared to combined radioimmunotherapy regimens (C12/aC4-iF and C12/aC4-oF). Only two genes were enriched based on p-value $<0.05$ and log2 fold-change $>0.5$. No genes were enriched upon radiomonotherapy.

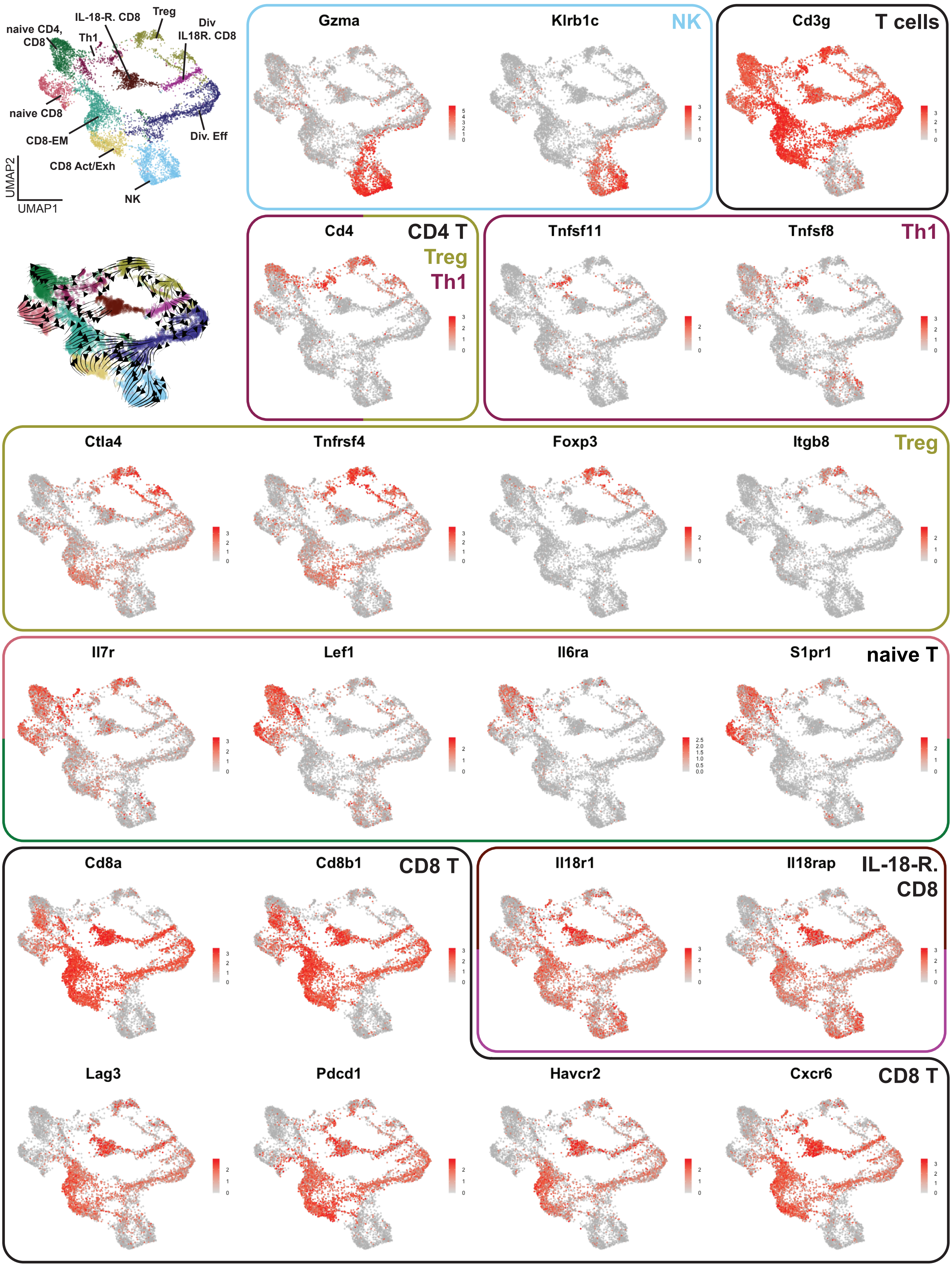

naive CD4, CD8
IL-18-R. CD8
Th1
Treg
Div IL18R. CD8
naive CD8
CD8-EM
Div. Eff
CD8 Act/Exh
NK
UMAP2
UMAP1
Gzma
Klrb1c
NK
Cd3g
T cells
Cd4
CD4 T
Treg
Th1
Tnfsf11
Tnfsf8
Th1
Ctla4
Tnfrsf4
Foxp3
Itgb8
Treg
Il7r
Lef1
Il6ra
S1pr1
naive T
Cd8a
Cd8b1
CD8 T
Il18r1
Il18rap
IL-18-R. CD8
Lag3
Pdcd1
Havcr2
Cxcr6
CD8 T

**Fig. S7. Marker genes for lymphoid subclustering.** UMAPs of lymphoid marker genes selected for subclustering (related to Fig. 5A and C). The two left top graphs show clustering and velocity as in Fig. 5A (upper graph) as well as clustering colored by treatment groups (lower graph).

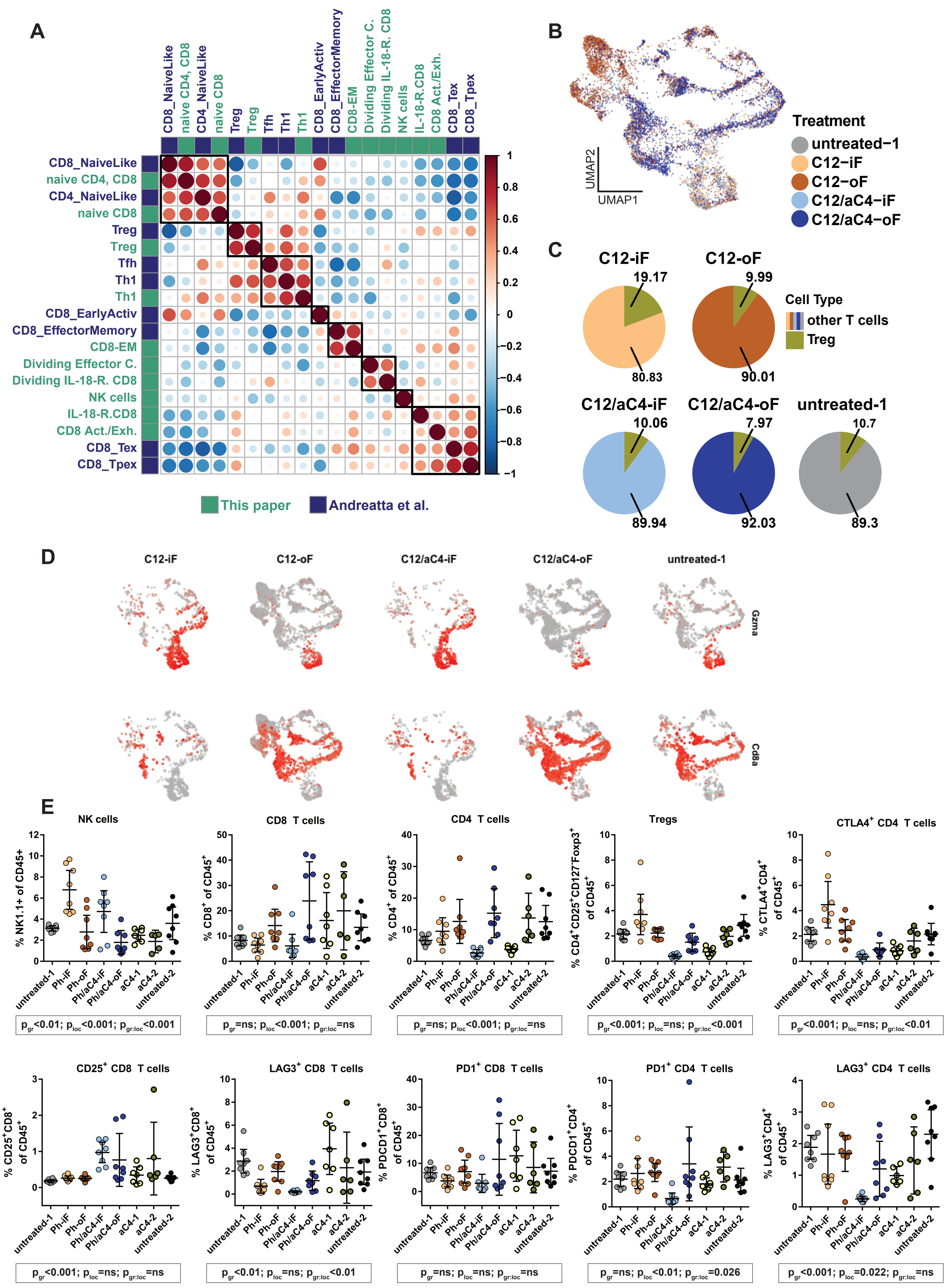

A
CD8_NaiveLike
naive CD4, CD8
CD4_NaiveLike
naive CD8
Treg
Treg
Tfh
Th1
Th1
CD8_EarlyActiv
CD8_EffectorMemory
CD8-EM
Dividing Effector C.
Dividing IL-18-R. CD8
NK cells
IL-18-R.CD8
CD8 Act./Exh.
CD8_Tex
CD8_Tpex
1
0.8
0.6
0.4
0.2
0
−0.2
−0.4
−0.6
−0.8
−1
This paper
Andreatta et al.
B
UMAP2
UMAP1
Treatment
untreated−1
C12−iF
C12−oF
C12/aC4−iF
C12/aC4−oF
C
C12-iF
19.17
80.83
C12-oF
9.99
90.01
Cell Type
other T cells
Treg
C12/aC4-iF
10.06
89.94
C12/aC4-oF
7.97
92.03
untreated-1
10.7
89.3
D
C12-iF
C12-oF
C12/aC4-iF
C12/aC4-oF
untreated-1
Gzma
Cd8a
E
NK cells
% NK1.1+ of CD45+
CD8 T cells
% CD8+ of CD45+
CD4 T cells
% CD4+ of CD45+
Tregs
% CD4+CD25+CD127-Foxp3+ of CD45+
CTLA4+ CD4 T cells
% CTLA4+CD4+ of CD45+
untreated-1
Ph-iF
Ph-oF
Ph/aC4-iF
Ph/aC4-oF
aC4-1
aC4-2
untreated-2
pgr<0.01; ploc<0.001; pgr:loc<0.001
pgr=ns; ploc<0.001; pgr:loc=ns
pgr=ns; ploc<0.001; pgr:loc=ns
pgr<0.001; ploc=ns; pgr:loc<0.001
pgr<0.001; ploc=ns; pgr:loc<0.01
CD25+ CD8 T cells
% CD25+CD8+ of CD45+
LAG3+ CD8 T cells
% LAG3+CD8+ of CD45+
PD1+ CD8 T cells
% PDCD1+CD8+ of CD45+
PD1+ CD4 T cells
% PDCD1+CD4+ of CD45+
LAG3+ CD4 T cells
% LAG3+CD4+ of CD45+
pgr<0.001; ploc=ns; pgr:loc=ns
pgr<0.01; ploc=ns; pgr:loc<0.01
pgr=ns; ploc=ns; pgr:loc=ns
pgr=ns; ploc<0.01; pgr:loc=0.026
pgr<0.001; ploc=0.022; pgr:loc=ns

**Fig. S8. Correlation to T cell reference confirms lymphoid subtypes.**

**(A)** Correlation heatmap of subclustered lymphoid clusters of this paper with a reference dataset (Andreatta *et al*, 2021). Modules of co-expressed genes were calculated on the reference datasets (see Materials and Methods) and gene expression was aggregated per cluster and module. Aggregated expression values were used for correlation. Color code and dot size show pairwise correlation from anti-correlation (-1, dark blue and large dot) over no correlation (0, white and small dot) to high correlation (1, red and large dot). Note that the reference did not include dividing cells and NK cells. This explains why Dividing Effector cells and Dividing IL-18-R CD8 show highest correlation to themselves, since like NK cells they do not have a counterpart in the reference. Gene modules as well as aggregated expression values can be found in **Table S8**.

(**B**) Lymphoid cells embedded in UMAP and color coded by treatment group reveals enrichment of naïve T cells in C12-oF tumors and of NK cells in C12-iF and C12/aC4-iF tumors, while C12/aC4-oF tumors are enriched in activated T cells.

(**C**) While Tregs were increased in oF tumors compared to all immune cells (Fig. 5B), due to an increase in T cells, the relative abundance of regulatory T cells among all T cells remains unchanged.

(**D**) Expression of NK cell marker Gzma and CD8 T cell marker Cd8a. Dividing lymphoid cells in iF tumors (C12-iF and C12/aC4-iF) are Gzma expressing NK cells, while in oF tumors (C12-oF and C12/aC4-oF) they are $CD8^+$ T cells.

**(E**) Flow cytometric analysis of the T cell / NK cell compartment in tumors following the treatment schedule depicted in Fig. 1A. Each dot represents an individual tumor.

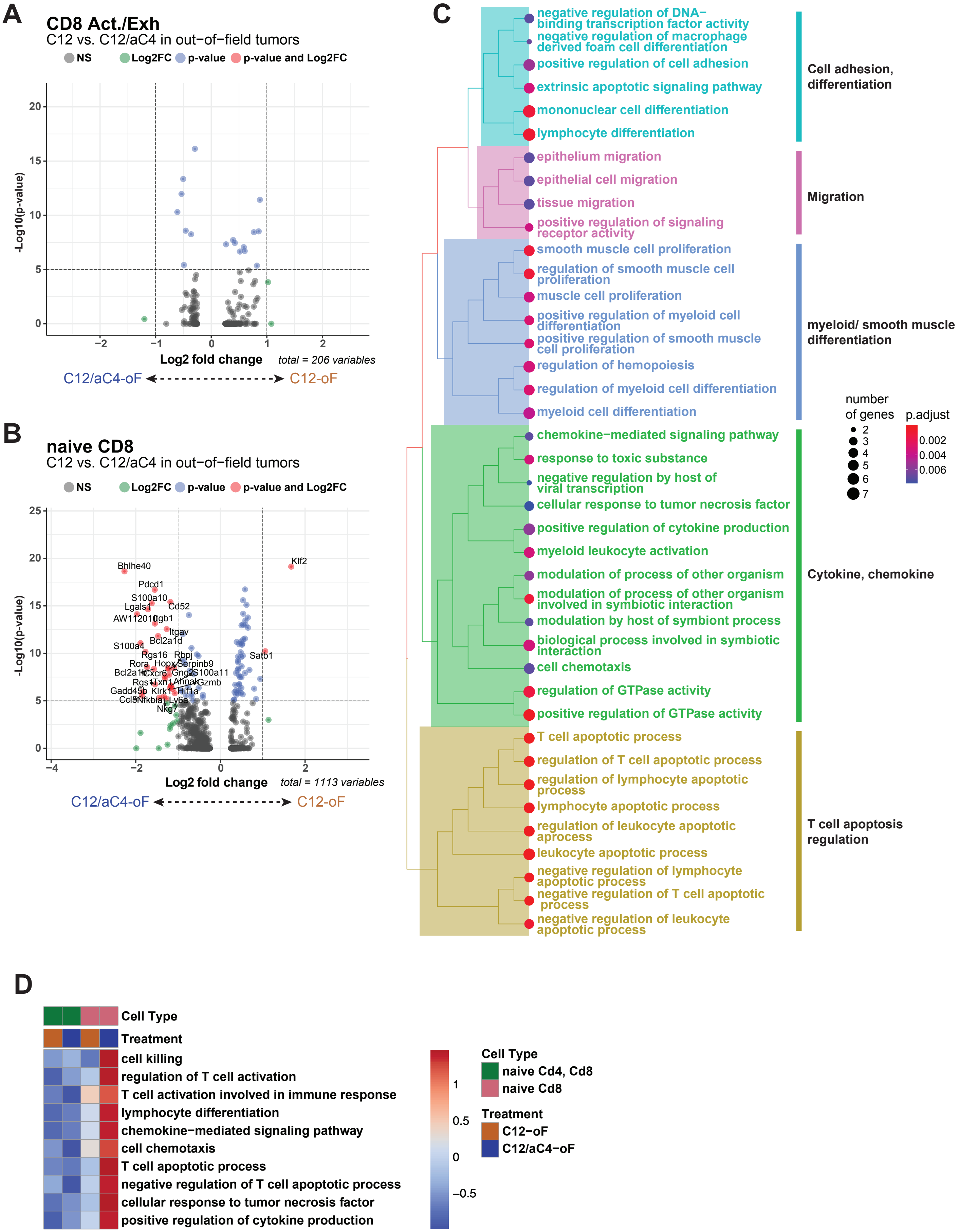
A
CD8 Act./Exh
C12 vs. C12/aC4 in out−of−field tumors
NS
Log2FC
p-value
p-value and Log2FC
-Log10(p-value)
Log2 fold change
total = 206 variables
C12/aC4-oF
C12-oF
B
naive CD8
C12 vs. C12/aC4 in out−of−field tumors
total = 1113 variables
Bhlhe40
Klf2
Pdcd1
S100a10
Lgals1
Cd52
Itgav
Bcl2a1d
S100a4
Rgs16
Rbpj
Satb1
Rora
Hopx
Serpinb9
Cxcr6
S100a11
Gzmb
Hif1a
Ly6a
Nkg7
C
negative regulation of DNA−binding transcription factor activity
negative regulation of macrophage derived foam cell differentiation
positive regulation of cell adhesion
extrinsic apoptotic signaling pathway
mononuclear cell differentiation
lymphocyte differentiation
epithelium migration
epithelial cell migration
tissue migration
positive regulation of signaling receptor activity
smooth muscle cell proliferation
regulation of smooth muscle cell proliferation
muscle cell proliferation
positive regulation of myeloid cell differentiation
positive regulation of smooth muscle cell proliferation
regulation of hemopoiesis
regulation of myeloid cell differentiation
myeloid cell differentiation
chemokine−mediated signaling pathway
response to toxic substance
negative regulation by host of viral transcription
cellular response to tumor necrosis factor
positive regulation of cytokine production
myeloid leukocyte activation
modulation of process of other organism
modulation of process of other organism involved in symbiotic interaction
modulation by host of symbiont process
biological process involved in symbiotic interaction
cell chemotaxis
regulation of GTPase activity
positive regulation of GTPase activity
T cell apoptotic process
regulation of T cell apoptotic process
regulation of lymphocyte apoptotic process
lymphocyte apoptotic process
regulation of leukocyte apoptotic aprocess
leukocyte apoptotic process
negative regulation of lymphocyte apoptotic process
negative regulation of T cell apoptotic process
negative regulation of leukocyte apoptotic process
Cell adhesion, differentiation
Migration
myeloid/ smooth muscle differentiation
Cytokine, chemokine
T cell apoptosis regulation
number of genes
p.adjust
D
Cell Type
Treatment
cell killing
regulation of T cell activation
T cell activation involved in immune response
lymphocyte differentiation
chemokine−mediated signaling pathway
cell chemotaxis
T cell apoptotic process
negative regulation of T cell apoptotic process
cellular response to tumor necrosis factor
positive regulation of cytokine production
naive Cd4, Cd8
naive Cd8
C12−oF
C12/aC4−oF

**Fig. S9. Combined radioimmunotherapy activates naïve T cells.**

**(A)** Analysis of differential gene expression in activated/exhausted $CD8^+$ T cells comparing C12/aC4-oF and C12-oF conditions. No differentially expressed genes were detected with p-value < 10^5 and log2 fold-change >1.

**(B)** Differential gene expression analysis in naïve $CD8^+$ T cells comparing C12/aC4-oF and C12-oF conditions. Combination strongly induced multiple genes with p-value < 10^5 and log2 fold-change >1. Differentially expressed genes for all clusters are reported in **Table S9**.

**(C)** Gene set enrichment analysis (GSEA) on genes enriched in C12/aC4-oF tumors as detected in (B). GO-terms are clustered according to similarity to derive functional groups (see Materials and Methods). Adjusted p-value is shown from blue to red and the number of genes contributing to the respective score are depicted by the size of the dot.

(**D**) Aggregated expression of selected enriched GO-terms of (C) per treatment group and cell type comparing naïve T cell clusters. Only the cluster naïve $CD8^+$ T cells showed induction of these pathways.

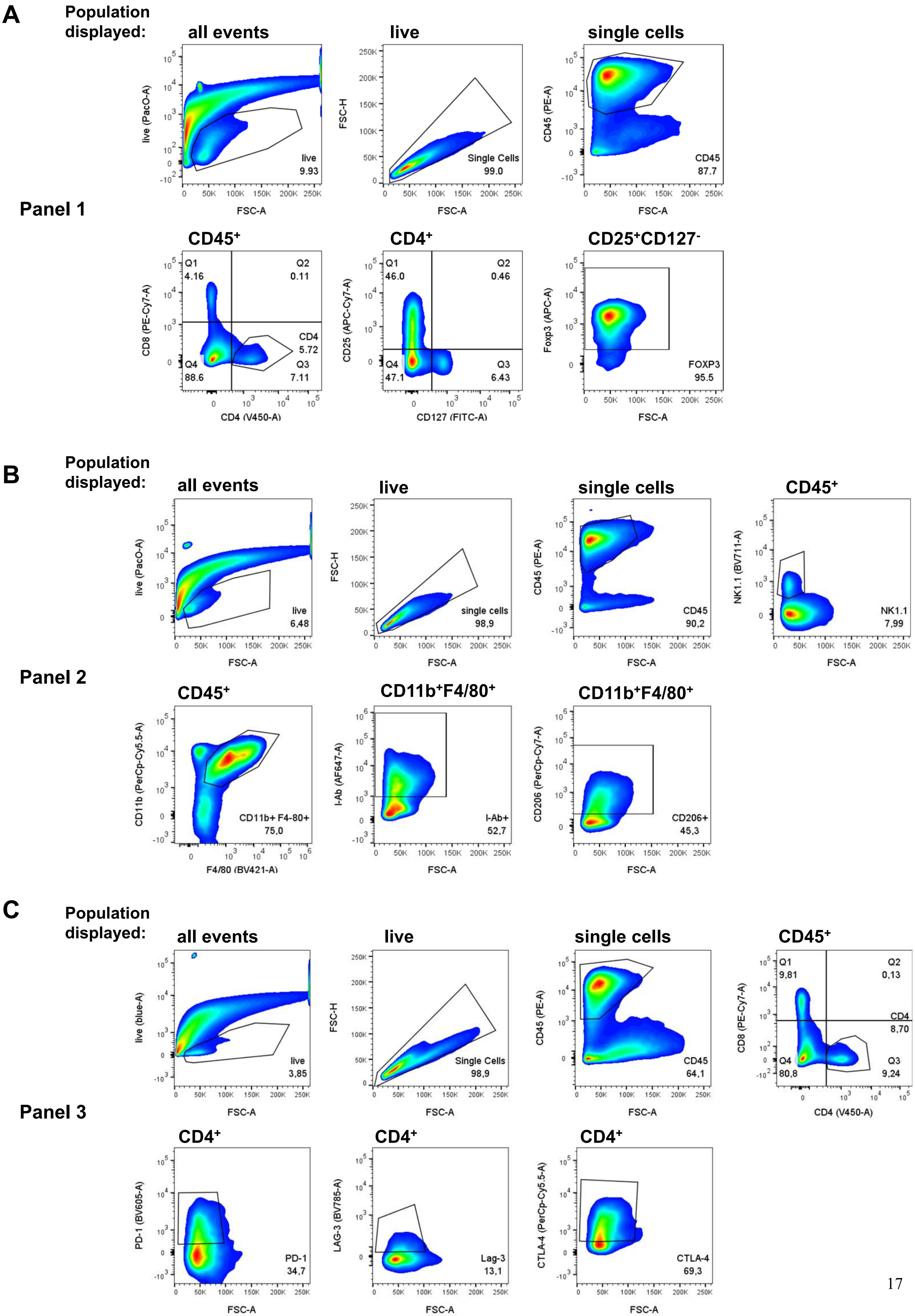
A
Population displayed:
all events
live
single cells
Panel 1
CD45+
CD4+
CD25+CD127-
live (PacO-A)
FSC-A
live
9.93
FSC-H
Single Cells
99.0
CD45 (PE-A)
CD45
87.7
CD8 (PE-Cy7-A)
CD4 (V450-A)
Q1
4.16
Q2
0.11
CD4
5.72
Q4
88.6
Q3
7.11
CD25 (APC-Cy7-A)
CD127 (FITC-A)
Q1
46.0
Q2
0.46
Q4
47.1
Q3
6.43
Foxp3 (APC-A)
FOXP3
95.5
B
Population displayed:
all events
live
single cells
CD45+
Panel 2
live
6,48
single cells
98,9
CD45
90,2
NK1.1 (BV711-A)
NK1.1
7,99
CD45+
CD11b+F4/80+
CD11b+F4/80+
CD11b (PerCp-Cy5.5-A)
F4/80 (BV421-A)
CD11b+ F4-80+
75,0
I-Ab (AF647-A)
I-Ab+
52,7
CD206 (PerCp-Cy7-A)
CD206+
45,3
C
Population displayed:
all events
live
single cells
CD45+
Panel 3
live (blue-A)
live
3,85
Single Cells
98,9
CD45
64,1
Q1
9,81
Q2
0,13
CD4
8,70
Q4
80,8
Q3
9,24
CD4+
CD4+
CD4+
PD-1 (BV605-A)
PD-1
34,7
LAG-3 (BV785-A)
Lag-3
13,1
CTLA-4 (PerCp-Cy5.5-A)
CTLA-4
69,3

**Fig. S10. Gating strategy for identification of immune cell populations.** Events were first gated on live cells, single cells and $CD45^+$ leukocytes.

(**A**) Panel 1: $CD45^+$ leukocytes were divided into $CD4^+$ and $CD8^+$ T cells. $CD4^+$ cells were further gated on $CD25^+CD127^-Foxp3^+$ cells to identify regulatory T cells.

(**B**) Panel 2: $CD45^+$ leukocytes were either gated on NK1.1 to identify NK cells or on $F4/80^+CD11b^+$ to identify tumor associated macrophages (TAMs). M1-like and M2-like TAMs were distinguished by I-$A^b$ and CD206 expression, respectively.

(**C**) Panel 3: $CD45^+$ leukocytes were divided into $CD4^+$ and $CD8^+$ T cells. Both $CD4^+$ and $CD8^+$ (not shown) were then analyzed for the surface expression of the immune checkpoint molecules PD-1, LAG-3, and CTLA-4.

**Table S1. Monoclonal antibodies and isotype controls used for flow cytometry.**

| | Specificity | Conjugate | Manufacturer | Clone | Clone of isotype control |
|---|---|---|---|---|---|
| **Panel 1** | CD45 | PE | BD | 30-F11 | - |
| | CD4 | V450 | BD | RM4-5 | R35-95 |
| | CD8 | PE/Cy7 | Biolegend | 53-6.7 | RTK2758 |
| | CD25 | APC/Cy7 | BD | PC61 | A110-1 |
| | CD127 | FITC | Biolegend | A7R34 | RTK2758 |
| | FoxP3 | APC | Invitrogen | FJK-16s | eBR2a |

| | Specificity | Conjugate | Manufacturer | Clone | Clone of isotype control |
|---|---|---|---|---|---|
| **Panel 2** | CD45 | PE | BD | 30-F11 | - |
| | CD11b | PerCP/Cy5.5 | Biolegend | M1/70 | RTK4530 |
| | F4/80 | BV421 | Biolegend | BM8 | RTK2758 |
| | I-Ab | AF647 | Biolegend | AF6-120.1 | MOPC-173 |
| | CD206 | PE/Cy7 | Biolegend | C068C2 | C068C2 |
| | NK1.1 | BV711 | Biolegend | PK136 | MOPC-173 |

| | Specificity | Conjugate | Manufacturer | Clone | Clone of isotype control |
|---|---|---|---|---|---|
| **Panel 3** | CD45 | PE | BD | 30-F11 | - |
| | CD4 | V450 | BD | RM4-5 | R35-95 |
| | CD8 | PE/Cy7 | Biolegend | 53-6.7 | RTK2758 |
| | PD-1 | BV605 | Biolegend | 29F.1A12 | RTK2758 |
| | LAG-3 | BV785 | Biolegend | C9B7W | RTK2071 |
| | CTLA4 | PerCP/Cy5.5 | Biolegend | UC10-4B9 | HTK888 |
